# Emergent properties of HNF4α-PPARγ network may drive consequent phenotypic plasticity in NAFLD

**DOI:** 10.1101/2020.02.18.953935

**Authors:** Sarthak Sahoo, Divyoj Singh, Priyanka Chakraborty, Mohit Kumar Jolly

## Abstract

Non-Alcoholic Fatty Liver Disease (NAFLD) is the most common form of chronic liver disease in adults and children. It is characterized by excessive accumulation of lipids in the hepatocytes of patients without any excess alcohol intake. With a global presence of 24% and limited therapeutic options, the disease burden of NAFLD is increasing. Thus, it becomes imperative to attempt to understand the dynamics of disease progression at a systems-level. Here, we decode the emergent dynamics of underlying gene regulatory networks that have been identified to drive the initiation and progression of NAFLD. We have developed a mathematical model to elucidate the dynamics of the HNF4α-PPARγ gene regulatory network. Our simulations reveal that this network can enable multiple co-existing phenotypes under certain biological conditions: an adipocyte, a hepatocyte, and a “hybrid” adipocyte-like state of the hepatocyte. These phenotypes may also switch among each other, thus enabling phenotypic plasticity and consequently leading to simultaneous deregulation of the levels of molecules that maintain a hepatic identity and/or facilitate a partial or complete acquisition of adipocytic traits. These predicted trends are supported by the analysis of clinical data, further substantiating the putative role of phenotypic plasticity in driving NAFLD. Our results unravel how the emergent dynamics of underlying regulatory networks can promote phenotypic plasticity, thereby propelling the clinically observed changes in gene expression often associated with NAFLD.

## INTRODUCTION

The dawn of the 21st century has seen a major global outbreak of obesity and metabolic syndrome. In 2016, the World Health Organization (WHO) estimated that nearly two billion people are either overweight or obese. Primary factors driving this non-communicable epidemic are caloric excess and a sedentary lifestyle, leading to chronic diseases [1]. In the liver, these chronic diseases manifest as Non-Alcoholic Fatty Liver Disease (NAFLD). It is the most common form of chronic liver disease among adults and children, with a global prevalence of 24% [1,2]. NAFLD is defined as an excessive accumulation of fat in the liver, with at least 5% of hepatocytes exhibiting accumulated triglycerides, without an excess intake of alcohol (daily intake <30 g in men and <20 g in women per day). The disease spectrum varies from benign hepatocellular simple steatosis (SS) of the liver characterized by excessive and abnormal retention of triglycerides and cholesterol esters to a more severe form, called Non-Alcoholic SteatoHepatitis (NASH), marked by chronic inflammation of the liver, possibly resulting in cell death and subsequent cirrhosis or fibrosis in the liver. In a small fraction of cases, it can progress to form hepatocellular carcinoma (HCC) [2].

NAFLD is a complex disease that is an intricate interplay of intertwined genetic and environmental factors [3–5]. On one hand, the Genome Wide Association Studies (GWAS) have identified variants in the genes PNPLA3, TM6SF2, GCKR, MBOAT7 and HSD17B13 that seem to be associated with susceptibility to and/or progression of NAFLD [6]. A single-nucleotide polymorphism in PNPLA3, causing I to M transition at position 148, remains to be the variant that is most robustly associated with the entire spectrum of NAFLD. The evidence for the heritability of NAFLD is strengthened by data derived from epidemiological, familial aggregation and twin studies [6]. On the other hand, the prevalence of NAFLD is strongly associated with metabolic syndromes (obesity, type 2 diabetes mellitus, insulin resistance, and dyslipidemia) [7]. Therefore, some recent attempts have even suggested renaming NAFLD as MAFLD (metabolic associated fatty liver disease) [8]. However, the association of NAFLD with obesity need not be universal; particularly, in Asian populations, NAFLD in the absence of obesity – so-called ‘lean-NAFLD’ – has been reported. Lean NAFLD patients may be characterized by different pathogenetic processes as compared to obese NAFLD ones [1,9–12].

Recently, a molecular-level understanding of NAFLD has been emerging through the identification of signaling pathways implicated in hepatic lipid homeostasis. MAPK, NF-kB, AMPK and AKT pathways among others have been identified to be dysregulated in NAFLD [13]. In NAFLD, the lipids accumulated in the liver are primarily derived from the serum fatty acid (FA) pool (60%), followed by a contribution from *de-novo lipogenesis* (DNL) (25%; 3-fold higher than healthy controls) and the remaining 15% from the dietary sources. Many genes involved in adipogenic programs play a key role in the initiation and progression of NAFLD [7]. Peroxisome proliferation-activated receptor gamma 2 (PPARγ), a master regulator of adipogenesis [14], is frequently upregulated in NAFLD [15]. Similarly, sterol regulatory binding element protein-1c, SREBP-1c (protein coded by the gene SREBF1), a major driver of hepatic DNL, is upregulated in NAFLD [7]. Consequently, many coregulators and downstream target genes of PPARγ and SREBP-1c are also perturbed in the context of NAFLD [15]. Another intriguing observation about NAFLD patients is their lower levels of hallmark liver tissue maintenance genes of the liver such as Hepatocyte Nuclear Factor 4α (HNF4α) and Hepatocyte Nuclear Factor 1α (HNF1α) [16–18]. HNF4α is an established master regulator of induction and maintenance of the hepatic cell state [19,20]. HNF4α knockout mice exhibit severe hepatomegaly (enlarged liver) and steatosis [16,19]. Similar to HNF4α-knockout mice, HNF1α knockout also leads to a fatty liver phenotype with increased fatty acid synthesis and steatosis in the liver [21].

The abovementioned studies identify various players associated with NAFLD in a correlative and/or causative manner. However, most studies focus on investigating how the upregulation or downregulation of one or a pair of these genes affect the final phenotype of the disease. Thus, the emergent dynamics of disease initiation and progression as a consequence of interactions among these genes in a regulatory network is not well understood. Consequently, it still remains elusive how these coordinated gene expression changes emerge as a result of the dynamics of underlying regulatory networks.

Here, we have identified a core regulatory network involving HNF4α, HNF1α, PPARγ and SREBP-1c, and show how interconnections among these key players can drive NAFLD. Our results highlight that this network can give rise to multi-stability, i.e. the co-existence of multiple phenotypes – hepatocyte state (high HNF4α, low PPARγ), adipocyte state (low HNF4α, high PPARγ), and a “hybrid” adipocyte-like state of the hepatocytes (high HNAF4a, high PPARγ). Our results show that during NAFLD, cells can switch their phenotype from being a hepatocyte to a hybrid adipocyte-like state and *vice versa*, thus controlling phenotypic plasticity in the context of NAFLD. The clinical data from NAFLD patients support the trends as predicted from the model, i.e. deregulated levels of HNF4α, HNF1α, PPARγ and SREBP-1c. These results offer important insights into the emergent systems-level dynamics of the regulatory network driving NAFLD.

## RESULTS

### Identification of a core HNF4α-PPARγ network in hepatocytes

As a first step, we gathered experimentally curated information about interconnections among HNF4α, HNF1α, PPARγ and SREBP-1c in the context of lipid homeostasis in the liver and its disruption during the progression of NAFLD. HNF4α and HNF1α were identified as among the six master regulators for human hepatocytes, using chromatoimmuno-precipitation and high-resolution promoter microarrays [22]. Both of them can activate each other as well as positively auto-regulate themselves [18,22]. Their expression levels are lower in fatty liver disease patients as compared to healthy controls [16]. HNF4α plays a crucial role in hepatic development as well as in hepatic lipid homeostasis. Mouse embryos deficient in HNF4α can initiate liver development but not transcriptionally activate the liver-specific genes [23]. Further, adult mice lacking hepatic HNF4α (Hnf4-LivKO) tend to have disrupted lipid homeostasis *in vivo* and display a fatty liver phenotype [24,25]. The liver of Hnf4-LivKO mice had higher levels of mRNA and protein of PPARγ as compared to the liver of wild-type mice [25]. Recent studies have revealed HNF1α as a direct transcriptional repressor of PPARγ in the context of hepatic steatosis [26].

PPARγ and SREBP-1c, on the other hand, are two master regulators of adipocytic cell-fate. PPARγ is necessary and sufficient for adipogenesis in mammals [14]; overexpression of PPARγ has been reported to induce adipogenesis [27]. SREBP-1c, the major transcriptional regulator of lipogenesis, induces the cohort of genes necessary for synthesizing fatty acids [28] and increases DNL in the liver, a hallmark of NAFLD [29]. SREBP-1c is induced during the differentiation of 3T3-L1 preadipocytes [30]. Both PPARγ and SREBP-1c and PPARγ are upregulated in obese patients with NAFLD [31]; they can both self-activate directly or indirectly [32–35] and augment each other [36–39]. SREBP-1c can transcriptionally activate PPARγ directly by binding to a putative E-box in the promoter of PPARγ [36] and increase the transcriptional activity of PPARγ indirectly via the production of ligands for PPARγ [37]. Similarly, SREBF1 is predicted to be a high confidence target of PPARγ [38]. Intriguingly, SREBP-1c has been reported to transcriptionally repress HNF4α *in vitro* and *in vivo* in the rodent liver [39]. It can also inhibit the transcriptional activity of HNF4α, suppressing the expression of hepatic gluconeogenic genes [40].

The abovementioned interactions put together results in a core regulatory circuit ***(FIG 1)*** that consists of two pairs of closely connected and self-activating transcription factors and a mutual inhibition between these two pairs. One pair (HNF4α and HNF1α) drives the hepatocytic cell state, while the other pair (PPARγ and SREBP-1c) drives the adipocyte cell state. A mutual inhibition between these pairs is reminiscent of a ‘toggle switch’ seen between many ‘master regulators’ of two (or more) divergent cell fates, as identified during multiple instances during embryonic development and disease progression [41,42].

**FIGURE 1:**
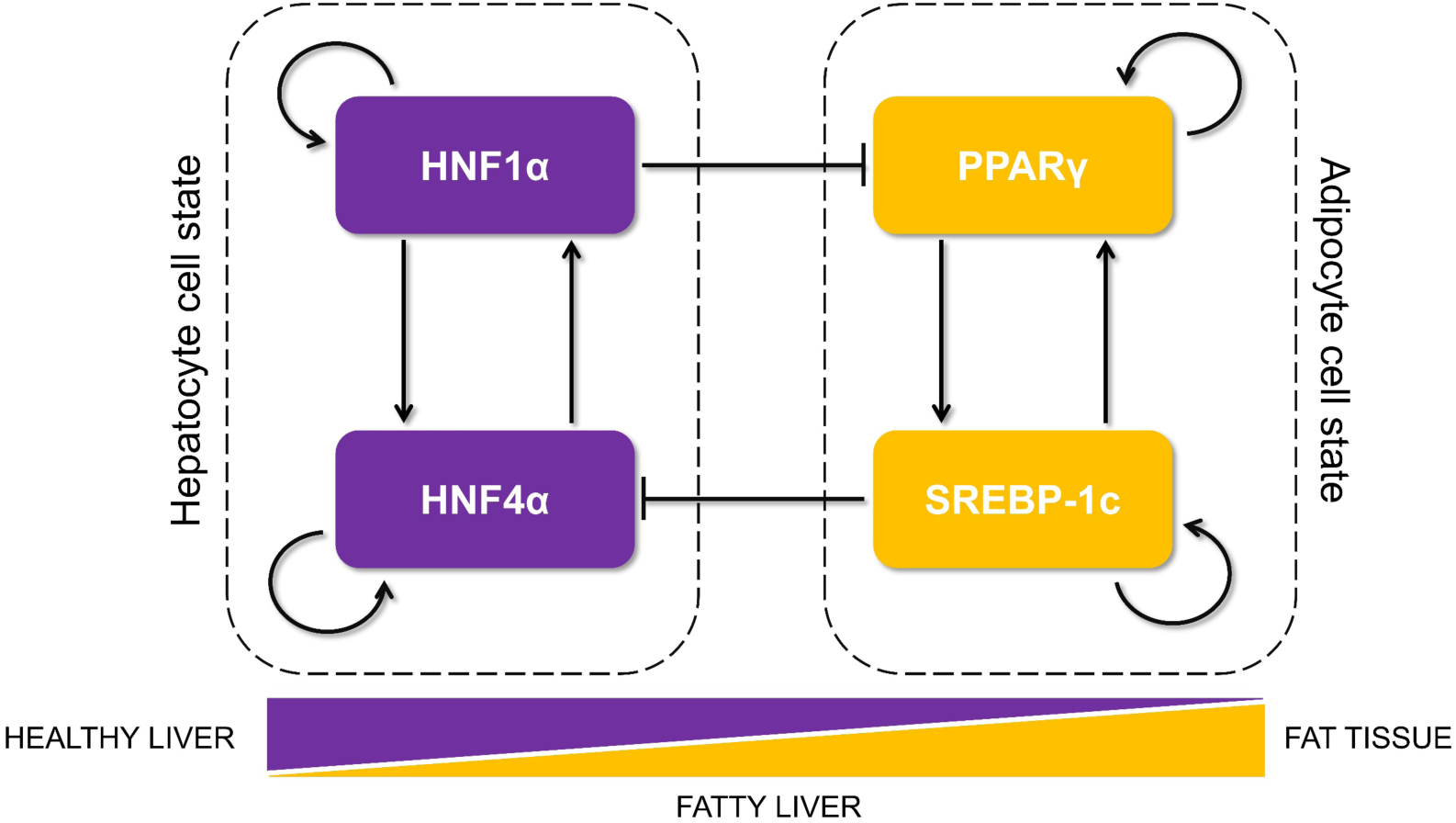
Proposed core regulatory network motif that controls lipid homeostasis in the liver and is dysregulated during the progression of NAFLD. The motif is composed of two tightly regulated, self-activated pairs of transcription factors (each pair shown in a dotted rectangle): HNF4α/ HNF1α (maintaining the hepatocyte cell state) and the PPARγ/SREBP-1c (driving an adipocytic cell state and implicated in initiation and progression of NAFLD). The arrows show transcriptional activation and the solid bars represent transcriptional repression. The relative levels of these proteins in a cell can determine its cell state as hepatocytes in healthy liver, adipocytes in the fat tissues or steatotic hepatocytes in livers of patients suffering from fatty liver.

### The emergent properties of this core regulatory network enable the existence of multiple phenotypes

Next, we investigated the dynamics emerging from this regulatory network. To characterize the robust dynamical properties of this network, we used a recently developed computational method - Random Circuit Perturbation (RACIPE) [43]. Given a topology of a regulatory network, RACIPE generates an ensemble of kinetic/biochemical models corresponding to the network topology, and later utilizes statistical tools to identify the robust dynamic properties emerging from the specific network topology. For each model in the ensemble, this method samples kinetic parameters from a biologically relevant chosen range of values. Thus, each kinetic model simulated via RACIPE takes a unique combination of parameters with the goal of capturing cell-to-cell heterogeneity in the biochemical reaction rates. An ensemble of these models thus denotes the behavior of a cell population, where each cell contains the given network but has different kinetic parameters from one another.

Here, each kinetic model is a set of four coupled ordinary differential equations (ODEs), each of which tracks the temporal changes in the levels of the four players (HNF4α, HNF1α, PPARγ and SREBP-1c) present in the core regulatory circuit. Each of these four has a rate of production and a rate of degradation; the production rate is governed by transcriptional regulation from other players (for instance, HNF4α is repressed by SREBP-1c), while the degradation term assumes first order kinetics and is independent of the levels of all the other genes). The set of differential equations are then solved numerically to attain the steady state values for each of the regulated players. These values are then mapped onto possible biological phenotypes represented as an attractor in the Waddington’s landscape [44] of cell fate determination ***(FIG 2A)***. For each given parameter set, depending on the initial condition, each of these molecular players can converge to one of the many possible steady states enabled by that parametric combination. Thus, the circuit considered here can be possibly multi-stable, potentially enabling phenotypic plasticity and/or heterogeneity.

**FIGURE 2:**
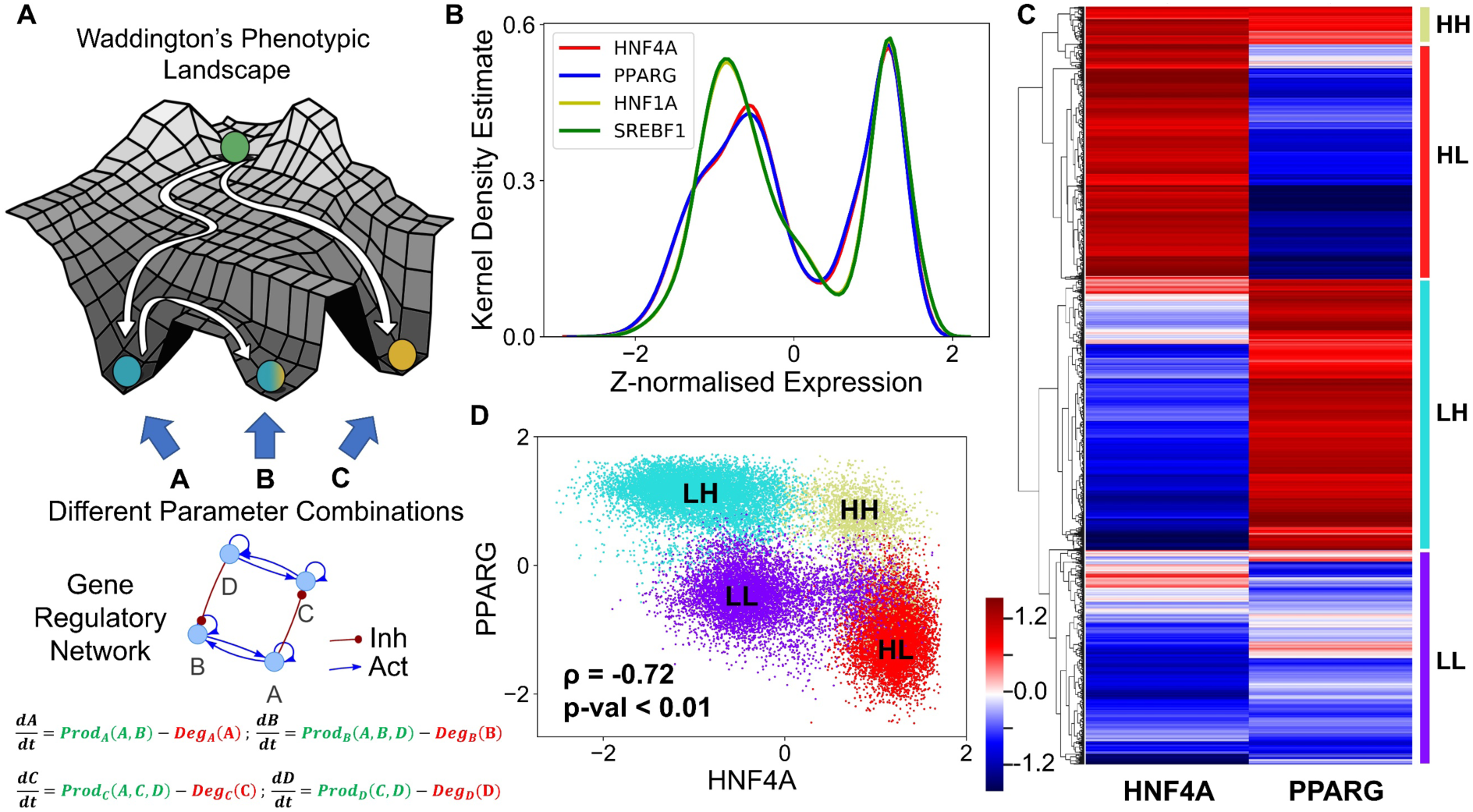
Core regulatory network may enable multiple phenotypes. **(A)** Schematic of Waddington’s phenotypic landscape highlighting the possibility of multiple states as a result of the underlying gene regulatory network; the dynamics of which can be modeled by a set of ordinary differential equations representing the interactions among them as shown below. **(B)** Kernel density plot showing the bimodal distribution of each component of the network, implying that each of the transcription factors (nodes) can exist in either a HIGH (H) or a LOW(L) state. The x-axis shows the z-normalized log2 expression values of the given component from RACIPE analysis. **(C)** Heatmap showing the relative levels of HNF4A and PPARG and the resulting four clusters. The color bar represents the relative levels of the individual components (z-normalized log2 expression values). **(D)** Scatter plot showing the existence of the four distinct clusters (defined based on hierarchical clustering) as the four states based on relative levels of HNF4A and PPARG (z-normalized log2 expression values). Spearman’s correlation was performed to obtain the correlation coefficient (ρ) and the corresponding p-value (p-val).

We collated the levels of HNF4α, HNF1α, PPARγ and SREBP-1c obtained from all parameter combinations and plotted the distribution of these levels. The distribution of levels of each of these four players is observed to be largely bimodal ***(FIG 2B, S1A)***, suggesting that each of these players can exist in either a “high” or a “low” state. Given that HNF4α and PPARγ are the key regulators of the hepatic and the adipocytic state respectively, we plotted the levels of HNF4α and PPARγ as a clustered heatmap. This analysis revealed four clusters corresponding to four different phenotypes: HNF4α-high and PPARγ-high (HH), HNF4α-low and PPARγ-low (LL), HNF4α-low and PPARγ-high (LH), HNF4α-high and PPARγ-low (HL) ***(FIG 2C)***. Interestingly, the HL and LH clusters were found to be the most abundant ***(FIG S2A)***, suggesting that this regulatory network can exist in two predominant states – HL (HNF4α-high, PPARγ-low) and LH (HNF4α-low, PPARγ-high). These states correspond to the hepatocyte and adipocyte phenotype respectively. Biologically speaking, this result is consistent with experimental observations that the exogenous overexpression of either HNF4α or PPARγ can drive the cell-fate to a hepatocyte (HL) or an adipocyte (LH) respectively [27,45] where the levels of the other master regulator is quite low ***(FIG S2B)***, i.e. either HNF4α/PPARγ ≫1 or HNF4α/PPARγ ≪1. Besides these two states, the network can exist in either a HH (HNF4α-high, PPARγ-high) or LL (HNF4α-low, PPARγ-low) state. The HH state can possibly be mapped onto the “hybrid” adipocyte-like phenotype of the hepatocytes where the levels of both HNF4α and PPARγ are high, similar to observations made in other scenarios where co-expression of mutually opposing master regulators have been reported [46,47]. The LL state can be thought of as an uncommitted “stem-like” phenotype where the levels of both master regulators are low and hence is not committed to either of the cell lineages. These observations remain qualitatively unchanged even when all the four regulatory players are considered for clustering ***(FIG S1G)***.

Consistent with the predominance of HL and LH phenotypes, the scatter plot of all steady state solutions revealed a significant negative correlation between HNF4α and PPARγ (***FIG 2D***), indicating that while cell-to-cell heterogeneity may enable varying levels of HNF4α and PPARγ in a cell population; these levels remain anti-correlated at a population level. Consistently, HNF4α levels correlated positively with those of HNF1α, but negatively with SREBP-1c, and PPARγ levels correlated positively with those of SREBP-1c, but negatively with HNF1α ***(FIG S1B-F)***. Put together, these results imply that the core regulatory circuit among HNF4α, HNF1α, PPARγ and SREBP-1c can give rise to multiple cell-fates in a biologically relevant parameter regime; with the more frequent states being a hepatocyte (HL - HNF4α-high, PPARγ-low)) and an adipocyte (LH - HNF4α-low, PPARγ-high).

### Multiple stable states (phenotypes) can co-exist, giving rise to phenotypic plasticity

Next, we investigated the possibility of whether two or more steady states (phenotypes) can co-exist, thus possibly enabling phenotypic plasticity. In the ensemble of parameters sets simulated via RACIPE; we found instances where this network leads to only one phenotype (mono-stable); as well as instances where it leads to two (bi-stability), three (tri-stability) or four (tetra-stability) phenotypes ***(FIG 3A)***. Multi-stability (i.e. the co-existence of more than one steady state) can enable cells to switch their phenotypes spontaneously (i.e. without the necessity of any strong external perturbation) but depending on the levels of intrinsic or extrinsic biological noise [48,49]. Interestingly, the percentage of parameter sets that gave rise to mono-stability was the least (***FIG 3B***), suggesting that this core regulatory network is more likely to be multi-stable and thus enable phenotypic plasticity in the context of NAFLD. Among the monostable solutions, HL and LH – corresponding to hepatocyte and adipocyte phenotype – were the most predominant ones ***(FIG 3C)***, reminiscent of our previous observations. Further inspection of the bi-and tri-stable solutions also revealed a similar trend. Among six (= number of ways to choose 2 out of 4 solutions) possible bi-stable phases (i.e. combinations of co-existing states), the most frequent phase was {HL, LH}, i.e. co-existence of HL and LH states ***(FIG 3D)***. The next two most frequent phases included either HL or LH as one of the states. Similarly, among the four possible tri-stable phases, the two more frequent ones contained both HL and LH, together with either LL or a HH one – {HL, LH, LL}, {HL, LH, HH} ***(FIG 3E)***.

**FIGURE 3:**
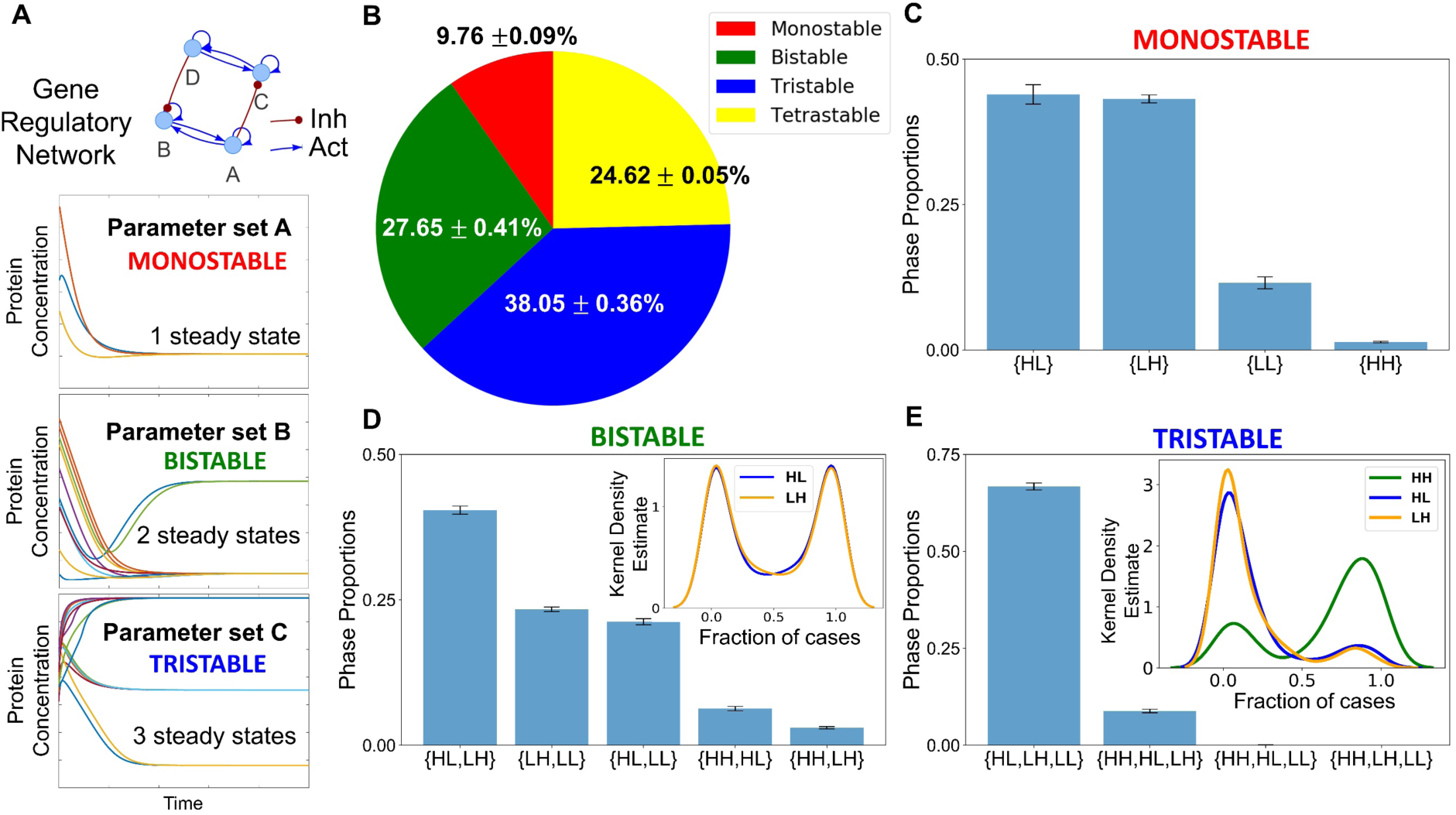
Co-existence and relative stability of multiple phenotypes, giving rise to multi-stable phases. **(A)** Schematic showing how a gene regulatory network for different parameter sets end up in a mono- or a multi- (bi-, tri-) stable regime. Depending on different initial conditions, the system (levels of the different regulators) can converge to one or more steady state levels, enabling mono- or multi-stability. **(B)** A pie chart showing that the fraction of parameter sets giving rise to mono-, bi-, tri-, tetra-stability. (mean ± standard deviation over three independent replicates through RACIPE analysis) **(C)** Bar plot showing the proportions of the various phases of the monostable solutions namely {HL}, {LH}, {LL} and {HH}. **(D)** Bar plot showing the proportions of the various phases possible for bi-stable solutions. The kernel density estimate plot in the inset shows the relative stability of the HL and LH states for the phase {HL, LH}. **(E)** Bar plot showing the proportions of phases possible for the tri-stable solutions. The kernel density estimate plot in the inset shows the relative stability of the HL, LH and HH states for the phase, {HL, LH, HH}. Error bars represent the standard deviation of the mean values of the phase proportions over three independent RACIPE replicates. Panels B-E are based on three replicates of RACIPE simulations on the network shown in Fig 1.

Within each phase, there may be varied stability of different co-existing phenotypes. Thus, we quantified the relative stability of the co-existing states in the different multi-stable phases. We calculated the fraction of randomly chosen initial conditions that converged to a particular state for all the parameter sets that enabled multi-stability (bi-, tri- or tetra-stability). Such calculation can provide insights into how likely the system will attain a particular state for an ensemble of randomly chosen initial conditions, and hence, can serve as a proxy measurement for the relative stability of that state in the phase. For instance, for a given parameter set corresponding to the phase {HL, LH}, depending on the sampling of initial conditions (say n=100), x of them can converge to HL state, while (100-x) converge to LH. We calculated the values of fractions of initial conditions leading to HL or LH and plotted the distribution of these values ***(FIG S3A-C)***. We found that both the HL and LH were equally likely to be attained if the system was left to start from a large set of randomly chosen initial levels of the different nodes present in the gene regulatory network when analyzed across parameter sets corresponding to {HL, LH} ***(FIG 3D inset)***. Such symmetry in the relative stability of states may emerge due to the symmetry in the network itself *(See the two sides of the network in* ***FIG 1***). For the phases {HH, LH} and {HH, HL}, the state HH seemed relatively more stable than either LH or HL; conversely, for the phases {HL, LL} and {LH, LL}, the state LL was found to be relatively less stable than HL or LL (***FIG S3D-G***).

The relative stability trends for the two tristable phases {HL, LH, LL} and {HL, HH, LH} and the tetrastable phase {HL, LH, LL, HH} reinforced our previous observations. LL was much less stable relative to HL or LH in the phase {HL, LH, LL} ***(FIG S3H)***, thus implying that this phase can be effectively considered to be equivalent to a bistable phase {HL, LH}. A possible interpretation would be that LL corresponds to a “stem-like” cell state that is inherently less stable and can quickly differentiate to an adipocyte or hepatocyte. Contrary to this, for the phase {HL, LH, HH}, the relative stability of the HH state was much larger than that of HL or LH states (***FIG 3E inset***). The HH state can be thought of as a “hybrid” adipocyte/hepatocyte state, similar to those observed in other tristable cell-fate decision networks [47,50,51]. This result, together with similar observations for the tetrastable phase ***(FIG S3I)***, emphasizes that a hybrid adipocyte-like state of hepatocytes (i.e. HH) may be prevalent in driving NAFLD.

Together, RACIPE analysis for this core regulatory network reveals a robust feature of this network topology – not only it allows cells to exist in more than one phenotype (HL, LH, HH), but also it can facilitate the stability of a HH state which may be quite relevant during the progression of NAFLD, i.e. a scenario where hepatocytes have not completely lost their identity but do simultaneously express various adipocytic markers and/or traits. The co-existence of HH with other states (HL, LH) raises the possibility that during the initiation and/or progression of NAFLD, hepatocytes may reversibly and dynamically switch to this adipocyte-like or hybrid adipocyte/hepatocyte phenotype, emphasizing the role of phenotypic plasticity in NAFLD.

After getting insights into robust dynamical properties of the core regulatory network across a range of parametric combinations, we studied what might possibly be driving NAFLD in a more realistic situation. NAFLD can be construed as a spectrum of diseases where hepatocytes can lose their hepatocytic identities to varying degrees and/or gain adipocytic identities to different extents, enabled by phenotypic plasticity of the underlying biological network (***FIG 4A***). Thus, the histopathological state of the liver seen in NAFLD is more likely to map onto the hybrid HH state as observed from the RACIPE results in which hepatocytes gain adipocytic characteristics without completely losing their hepatocytic identity. Thus, we hypothesized that NAFLD may progress via a switch from the HL (hepatocyte) state to the HH state (hybrid or adipocyte-like state of hepatocyte).

**FIGURE 4:**
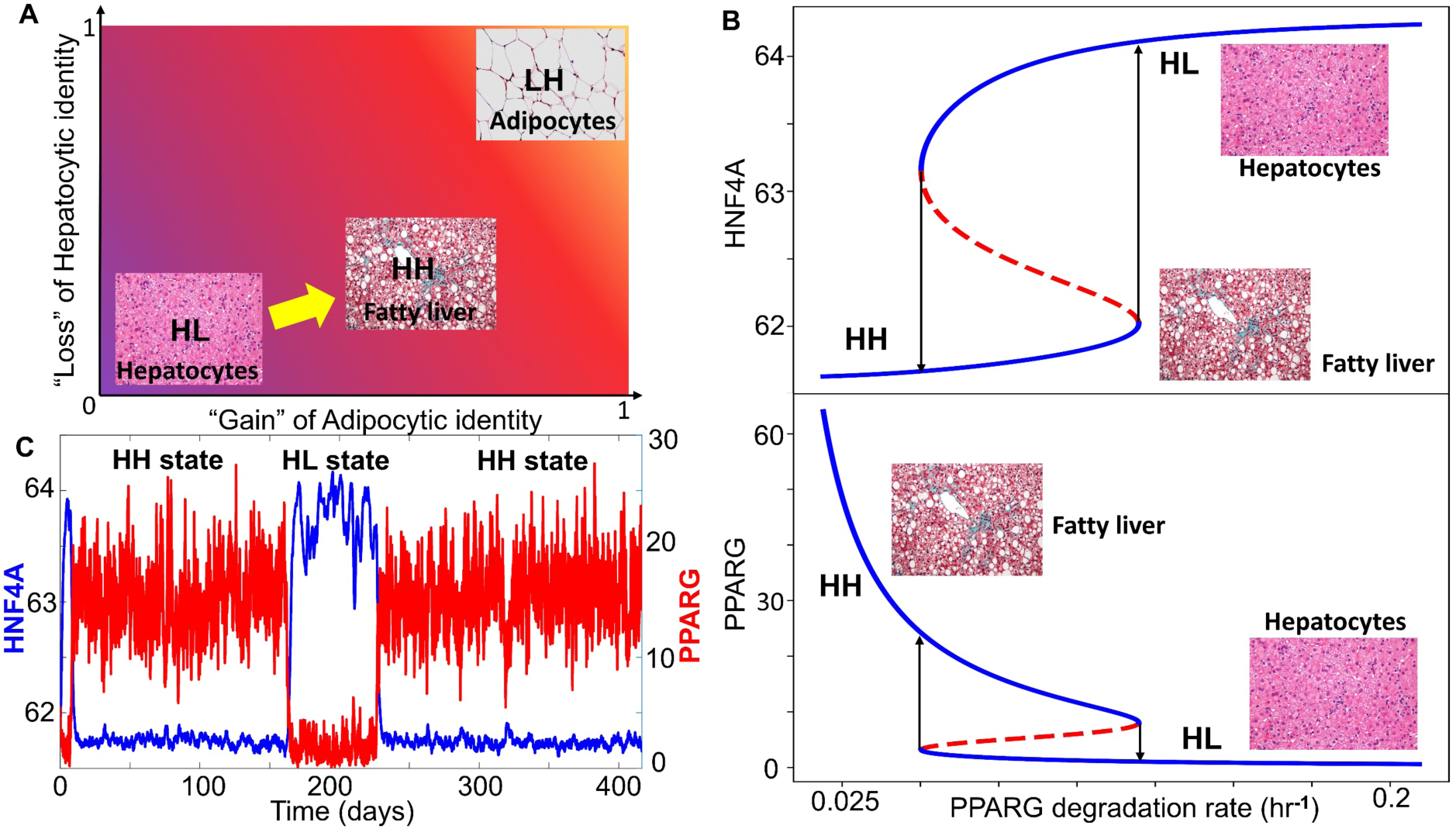
NAFLD as a bistable system involving a phenotypic switch from a hepatocyte to a “hybrid” adipocyte-like phenotype of the hepatocytes. **(A)** Schematic showing the phenotypic transition that may happen in the context of NAFLD where the disease can be highly diverse in terms of the extent of loss of hepatocytic and/or gain of adipocytic characteristics. **(B)** Bifurcation plots showing the phenotypic transitions, i.e. switch in the levels of HNF4A (top panel) and PPARG (bottom panel) when the degradation rate of PPARG is used as a bifurcation parameter. Solid blue lines denote the stable states, dotted red lines denote the unstable states of the system. **(C)** Stochastic simulations of the core gene regulatory network showing switching in HNF4A and PPARG levels (degradation rate of PPARG = 0.08 hr^-1^). (Image credit for ‘hepatocyte’, ‘fatty liver’, ‘adipocytes’ in A and B: Wikimedia Commons).

We estimated parameters ***(See methods)*** from relevant experimental literature to calibrate our model and identify whether a phenotypic switch is implicated during NAFLD. During NAFLD, the upregulation of Hsp90 levels can suppress the degradation of PPARγ, thus increasing PPARγ signaling [52]. We examined the effect of decreasing the degradation rates of PPARγ. Thus, as the degradation rate of PPARγ decreases, the steady state levels of PPARγ, which are generally low in the hepatocytes, increase constantly which switch beyond a certain threshold in PPARγ ***(FIG 4B bottom panel)***. As the degradation rate of PPARγ is varied, HNF4α levels drop modestly ***(FIG 4B top panel)*** indicating that the hepatocytic identity is not lost completely. Thus, at a higher degradation rate of PPARγ, the cell is in a hepatocytic state (solid blue line corresponding to ‘HL’); while lower degradation rates of PPARγ enables a switch to hybrid state (solid blue line corresponding to ‘HH’) ***(FIG 4B)***. This process can be viewed as “trans-differentiation” of the cell, i.e. conversion of hepatocytes to hybrid adipocyte-like cell state by simultaneous repression of the original tissue (liver) homeostatic mechanisms and activation of a different tissue-specific (adipose tissue) program. This result is consistent with observations of multiple adipocytic markers reported in histological samples of NAFLD patients [53]. In the liver, such trans-differentiation has also been observed for quiescent hepatic stellate cells that can attain adipogenic or myogenic characteristics depending on the relative abundance of adipogenic or myogenic genes respectively [54].

We further examined whether this core regulatory network can enable phenotypic switching under the influence of biological noise [55]. To mimic various sources of biological noise, we simulated the system stochastically *(see* ***Materials and Methods*** *for details)* and observed fluctuations in the levels of both PPARγ and HNF4α. Sometimes, these fluctuations were large enough to trigger a transition from the HL (hepatocyte; HNF4α high-PPARγ low) state to the HH state (hybrid; HNF4α high-PPARγ high) and *vice versa* ***(FIG 4C)***. The stability of the HH state can be assessed by the observation that the system may tend to spend a relatively long time in the HH state rather than the HL state. It was encouraging to note that the timescale of switching is in the order of three months, a timescale which corroborates excellently with clinical observations that NAFLD patients who adopt aggressive lifestyle changes may reverse NAFLD to a significant degree in one month [56]. Overall, we find that in the biologically relevant range of parameters under which NAFLD seems to be operating, hepatocytes may undergo a phenotypic switch to a hybrid adipocyte-like state of hepatocyte by partially activating the adipogenic program.

### The topology of the core regulatory network is designed to enhance phenotypic plasticity

Next, we investigated how unique are these observed traits such as multi-stability to the topology of the core network underlying NAFLD. In other words, we asked what the salient features of the topology of core regulatory network formed by reported interconnections among HNF4α, HNF1α, PPARγ and SREBP-1c are ***(FIG 1)***. To discern the effect of network topology, we created an ensemble of randomized networks, where we swapped/shuffled many links in the network with one another while maintaining the number of links that are incident upon and emanate from all individual nodes in the network. Such randomization enables us to dissect the contribution of network topology to the above-mentioned network features. For the core regulatory network shown in ***FIG 1***, 44 such “hypothetical” cases are possible (see **Methods**). One such example of a “hypothetical” network has been shown in ***FIG* 5A *inset***, where two activatory and two inhibitory links have been shuffled with respect to the “wild type” core network topology (***FIG 1***). In the “wild type” network, HNF1α inhibits PPARγ and SREBP-1c inhibits HNF4α, while in this “hypothetical network”, HNF1α activates PPARγ and SREBP-1c activates HNF4α. Moreover, in the “wild type” network, HNF1α activates HNF4α and PPARγ activates SREBP-1c, but in this “hypothetical” network, HNF1α inhibits HNF4α and PPARγ inhibits SREBP-1c.

_First, we calculated the state frequency of each of the 44 “hypothetical” (i.e. randomized) networks. We then computed the Jensen-Shannon distance (JSD) of the state frequency distribution of each of these “hypothetical” randomized networks from the observed state frequency of the “wild type” (i.e. core) regulatory network. JSD is a measure of distance between two distributions and varies between 0 and 1 [57]. JSD=0 implies that the two distributions are identical, while JSD=1 implies that the distributions are completely non-overlapping ***(FIG S6A)***. Upon plotting the distribution of the JSD of the 44 “hypothetical” networks, we found that none of the distributions were close to the observed distribution of the “wild type” regulatory network **(*FIG* 5A, S4A, S4C)** (minimum value of JSD = 0.22). This result signifies that the phenotypic distribution attained from the core regulatory network is unique to that network topology.

Next, we quantified the plasticity of the “wild type” network and 44 “hypothetical” networks. The plasticity of a network is defined to be the fraction of parameter sets that enable multi-stable solutions, out of the total number of parameter sets considered (n=10,000 here). We found the “wild type” network to possess the maximum plasticity as compared to each of the 44 “hypothetical” networks **(*FIG 5B*, S4B, S4D)**, highlighting that the specific network topology among these master regulators may be optimized to enable maximum plasticity.

Self-activation loops are known to contribute largely to the plasticity in many biological systems [58]; thus, we quantified the contribution of different self-activation loops on plasticity. The core network contained four self-activation loops: each for HNF4α, HNF1α, PPARγ and SREBP-1c. We created an ensemble of models that covered all possible combinations of self-activation loops: only one, two, or three out of the four nodes have such self-activation loops. We observed that the more the number of self-activation loops, the higher the plasticity score ***(FIG 5C)***. Put together, these results underscore that the topology of the core regulatory network plays crucial roles in enabling phenotypic plasticity during NAFLD progression.

**FIGURE 5:**
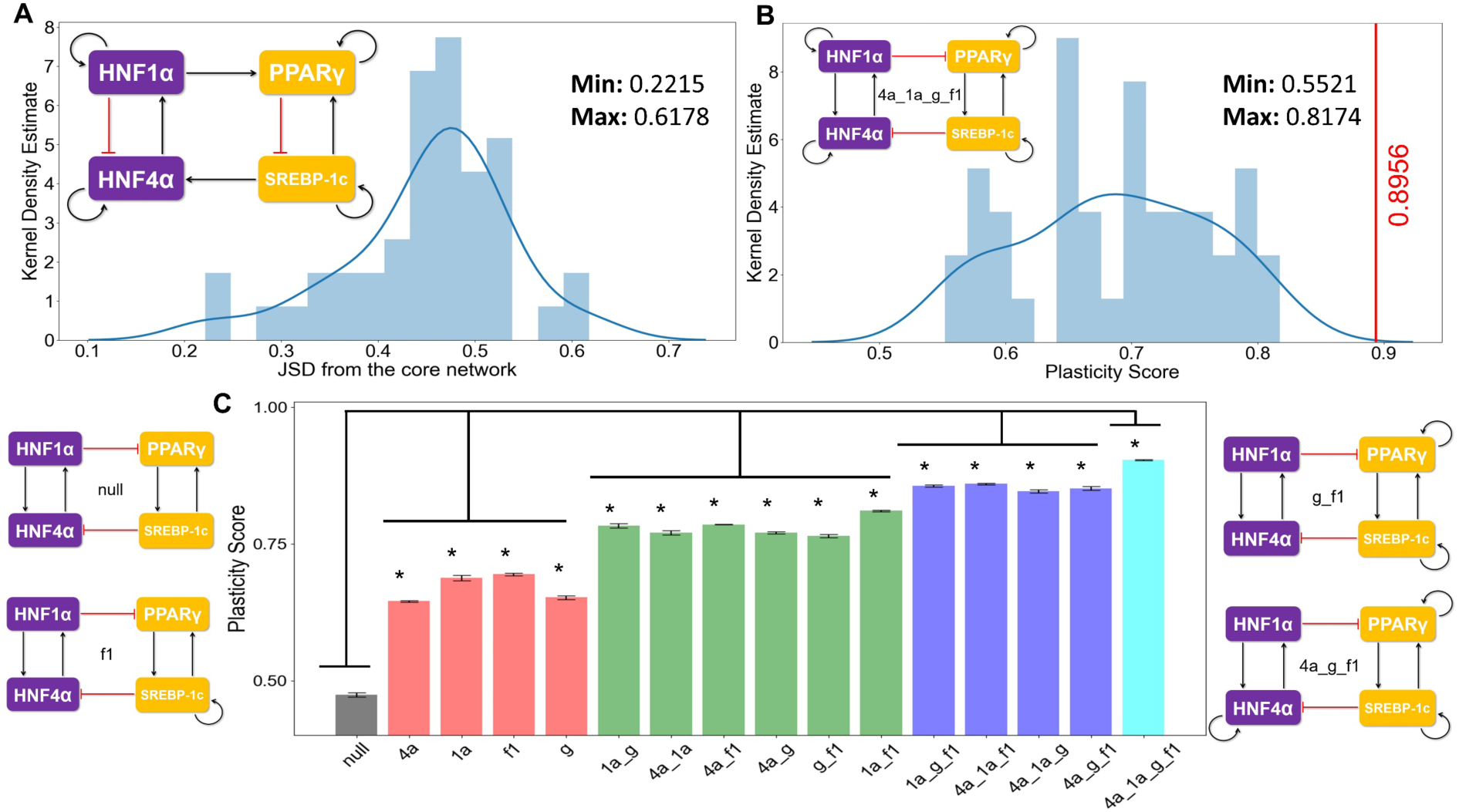
Proposed Core regulatory module for NAFLD enables maximum plasticity. **(A)** Kernel density estimate plot with the histogram overlaying showing the distribution of the Jensen-Shannon Distance (JSD) between the frequency of state distribution of the “hypothetical” randomized networks from the “wild type” core regulatory network. The maximum and minimum values of the JSD observed are mentioned in the top right side in the figure. An example of a randomized network is shown at the top left corner. **(B)** Kernel density estimate plot with the overlaying histogram showing the distribution of the plasticity scores of the various random networks. The core regulatory circuit (codenamed 4a_1a_g_f1) is shown in the inset. The red vertical line indicates the plasticity score (=0.89) of the “wild type” core regulatory circuit which is greater than all other “hypothetical” randomized networks. The maximum and minimum values of the JSD observed are mentioned in the top right side in the figure. **(C)** Bar plots showing the plasticity scores of the modified core regulatory networks with a varying number of direct self-activation loops. A few instances of the modified networks and the corresponding codenames are drawn alongside (null indicates no self-activation loops; 4a, 1a, g, f1 denotes the network topology having self-activations on HNF4A, HNF1A, PPARG and SREBF1 respectively). (* indicates a significant difference of the plasticity scores (p-val < 0.05) when compared via the student’s t-test with the plasticity of the null network)

### Clinical data support the model predictions

Finally, we tested whether our model predictions about phenotypic plasticity in NAFLD are consistent with the available clinical data. One key prediction of the model was that the expression levels of HNF4α and PPARγ should be negatively correlated ***(FIG 2D)***. We observed such trends in multiple clinical datasets (GSE66676, GSE33814, GSE37031), corresponding to NASH and/or NAFLD patients that showed PPARγ and HNF4α to be negatively correlated ***(FIG 6A)***. These results indicated that the core programs of hepatocytic and adipocytic identity are inversely correlated. As expected, the levels of HNF4α and HNF1α were positively correlated ***(FIG S5A)*** while that of the HNF1α and PPARγ were negatively correlated ***(FIG S5B)***.

**FIGURE 6:**
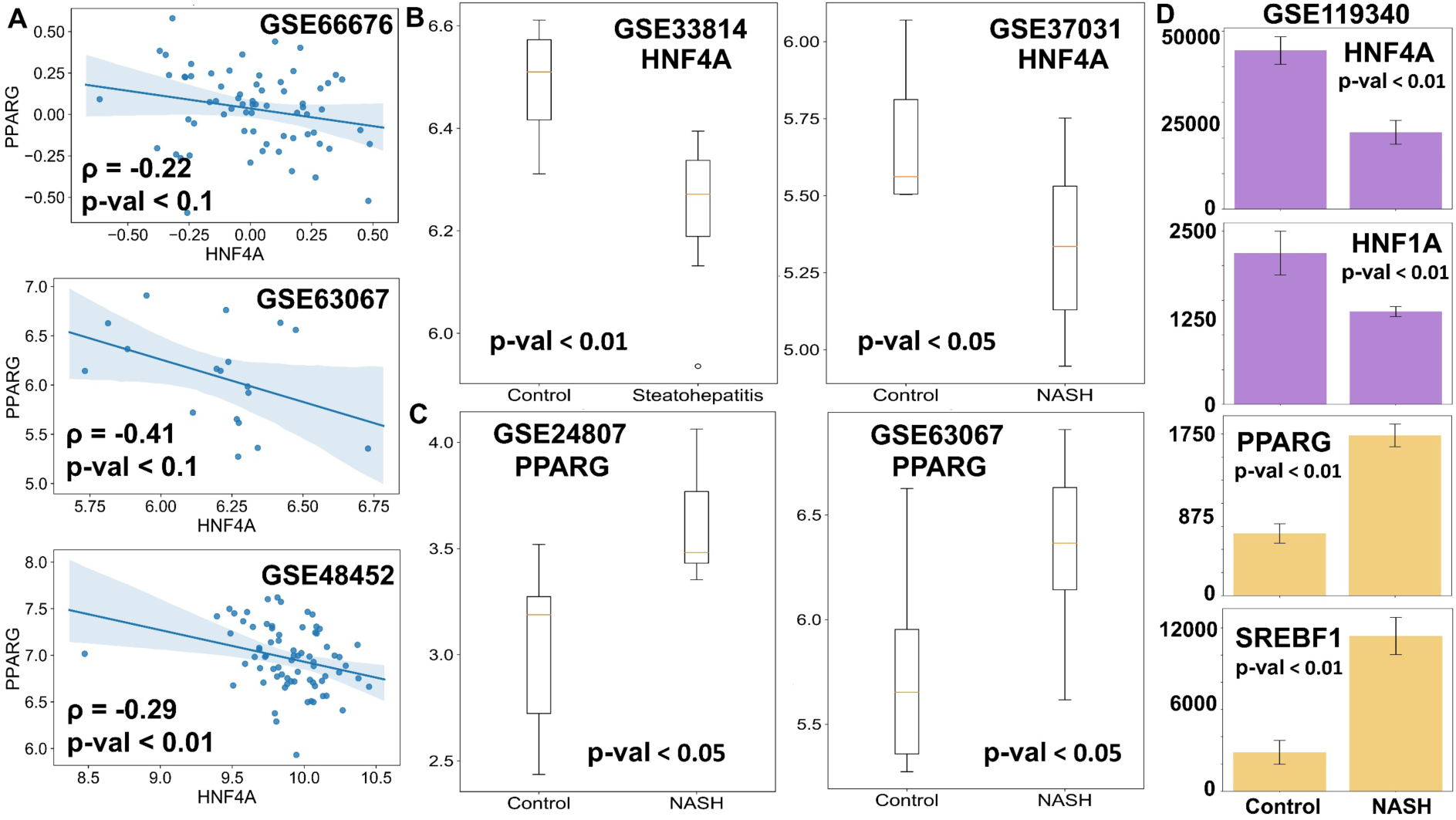
Empirical evidence supports the model predictions. **(A)** Scatter plots between the expression levels of HNF4A and PPARG in clinical samples. Spearman correlation coefficient is given by ρ, and p-val denotes the corresponding p-value. **(B, C**) Comparison of mRNA levels (log2 normalized) of HNF4A (B) and PPARG (C) in the liver of NASH patients and healthy controls. **(D)** Comparison of levels of HNF4A, HNF1A, PPARG and SREBF1 in mouse liver for control case vs. mouse model of NASH. Expression values are listed as TPM (transcripts per million) for given RNA-seq data. p-value (p-val) given for student’s t-test in B-D.

Next, we examined whether the HNF4α levels are significantly downregulated and/or PPARγ levels are upregulated in the case of NASH, a more severe stage of fatty liver disease. Indeed, HNF4α was found to be higher in the case of the normal liver than in the case of NASH livers ***(FIG 6B)***. Similarly, PPARγ levels were found to be amplified in livers of NASH patients in comparison to healthy livers ***(FIG 6C)***. These observations point out that NAFLD progresses via the simultaneous suppression of the hepatic program controlled by HNF4α and activation of the adipogenic program controlled primarily by PPARγ. Consistently, in mouse models of NASH, the liver showed downregulation of HNF4α and HNF1α and up-regulation of PPARγ and SREBP-1c, as compared to controls ***(FIG 6D)***. Intriguingly, this pattern was observed for hepatocytes, but not in liver endothelial cells, suggesting that this regulatory circuit may be specifically operative in hepatocytes, at least in mouse models ***(FIG S5C)***. Put together, these clinical observations strongly support that the proposed core gene regulatory network may underpin phenotypic plasticity in the context of NAFLD initiation and progression.

## DISCUSSION

With the rise of obesity and sedentary lifestyle-induced metabolic disorders, diseases such as NAFLD are on the rise with a very limited number of treatment options available which are not proven to be very effective [59]. The world has seen a drastic increase in the prevalence of NAFLD in the 21^st^ century and the number of cases continue to rise. The global incidence of NAFLD and its specific histological phenotype, NASH, have risen dramatically (15% in 2005 to 25% in 2010, and 33% in 2005 to 59.1% in 2010 respectively) [1]. Thus, it becomes imperative to study the initiation and progression of NAFLD through its various phenotypic stages to devise efficient and robust intervention measures to control the spread of this epidemic. Multiple studies have shown the importance of various key regulators and genetic alterations for the development and progression of NAFLD [60,61]. However, there are only a few attempts to analyze NAFLD from a systems biology perspective, i.e. decoding how these different regulators interact in the context of NAFLD [17,62–65]. Thus, mechanism-based modeling studies are required to elucidate these mechanisms form a dynamical systems perspective.

Here, we identified and computationally modeled a core gene regulatory network that appears to be crucial to explain various features of NAFLD. Specifically, we modeled the interactions among two hallmark liver homeostatic transcription factors, HNF4α and HNF1α, and two master regulators of the adipocytic cell fate and lipid homeostasis respectively, PPARγ and SREBP-1c. Our results show that the dynamical interactions between these transcription factors can drive a hepatocytic or an adipocytic cell fate program in a cell. This core regulatory network is also capable of existing in a “hybrid” state, adipocyte like phenotype of the hepatocytes, which might correspond to observations during the progression of NAFLD. This “hybrid” state is characterized by the presence of large lipid droplets in the hepatocytes with an increased expression of adipocytic makers and enhanced release of various pro-inflammatory cytokines such as IL-6, IL-18 and TNFα, both of which are hallmark features of adipocytes [53]. We also showed that the hepatocytes can spontaneously switch to form the “hybrid” state under the influence of biological noise. Further, we show that this core gene regulatory network exhibits higher levels of plasticity both due to its topology and the abundance of self-activation loops. Self-activation is frequently seen in biological networks, and its combination with mutually inhibitory circuits, as observed in the context of NAFLD, has been reported to amplify the likelihood of multi-stability and consequent phenotypic plasticity [42,58]. Thus, targeting the interactions that drive phenotypic plasticity and thus reducing the frequency of switching between hepatocytic and the “hybrid” states can be thought of as a potential therapeutic strategy for NAFLD.

The predictions of our model are also supported by clinical data of patients suffering from NAFLD. Overall, we can construe that the phenotype seen in NAFLD is due to a “trans-differentiation” process where the cells can switch from a stable attractor state (in this case, the hepatocytes) to another one (in this case, an adipocyte-like “hybrid” state of hepatocytes) ***(FIG 7)***. While further work needs to be carried out to elucidate the existence and extent of phenotypic switching in NAFLD patients, systems biology approaches similar to those presented here can have potential translational implications in terms of sub-phenotyping of patients and guiding subsequent drug repurposing and development [66,67]. For instance, the gut microbiota of lean NAFLD patients is strikingly different from that of obese NAFLD patients; such diverse microenvironmental traits may alter the interactome landscape in NAFLD pathogenesis [68].

**FIGURE 7:**
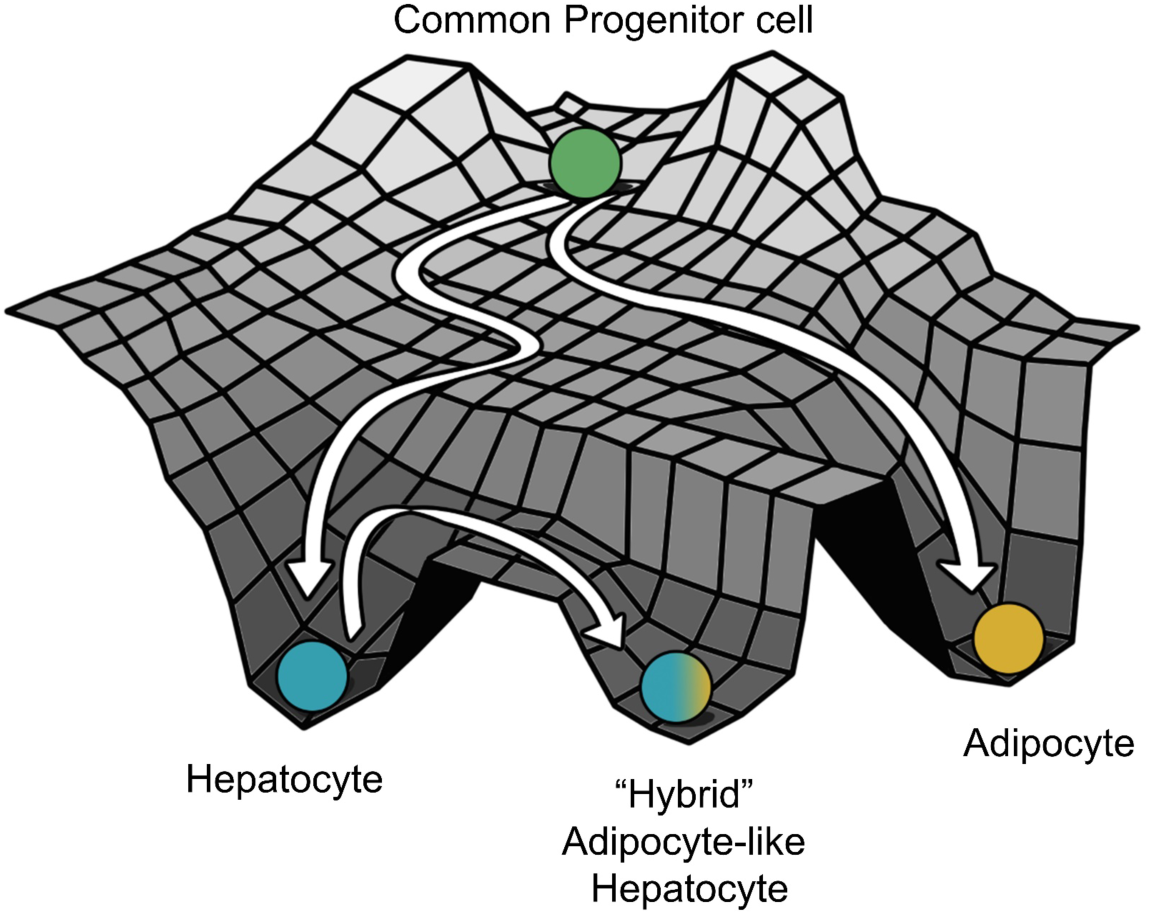
A schematic showing hepatocytes in a fatty liver as a distinct cell state due to “trans-differentiation” form the normal hepatocytes to a much more adipocyte like “hybrid” cell state on the Waddington’s landscape (The Strategy of the Genes, Waddington C, 1957).

Multistability is a hallmark of many cell-fate decision networks [69]. Multistable systems often display hysteresis; in other words, an asymmetry in the trajectory of ‘forward’ vs. ‘backward’ responses [70,71]. To establish the relevance of multistability in context of NAFLD, a proposed *in vitro* experiment would be to catalog the changes in levels of various adipocytic and/or hepatocytic markers at a single-cell scale and in a dose- and/or time-dependent manner, when hepatocytes are exposed to fatty acids and triglycerides driving NAFLD [72]. Another feature of multistable systems can be ‘spontaneous switching’ among phenotypes under the influence of biological noise [73]. To test this feature, one can isolate a population of steatotic hepatocytes and observe if it can give rise to non-steatotic hepatocytes (implying reversibility) when cultured separately *in vitro*. The extent of reversibility can depend on various factors such as remodeling of the cellular microenvironment and/or epigenetics [74]. Thus, it would also be interesting to quantify the reversibility of a fatty liver phenotype as the disease progresses from NAFLD to a more inflammatory state, NASH [75]. Specifically, it will be intriguing to examine how various players that are upregulated due to initial phenotypic transition (PPARγ and SREBP-1c) are stabilized due to the activation of the immune response, which is known to be a hallmark of NASH [75,76]. Another important trait that can impact the dynamics of NAFLD is the fluctuations of molecules during the fasting-feeding cycle [77]; for instance, SREBP-1c levels can increase drastically upon fasted mice being refed [78]. It should be noted that although our model offers a possible explanation for NAFLD progression, such phenotypic switching among adipocytes and hepatocytes has not been yet reported in adult homeostatic conditions, given their different developmental lineages, with adipocytes generally believed to be arising from the mesoderm [79] while the hepatocytes arising from the endoderm [80], although functional hepatocytes have been reported to be generated from human adipose-derived stem cells [81].

There are various limitations of the computational model considered here. First, the network proposed here is, by no means, exhaustive; various other mechanisms may alter the dynamics of the regulatory network and consequent relative stability and phenotypic switching. Second, the effect of various genetic variants has not been incorporated into the model at this first step. Third, our current model considers largely transcriptional factors, while both microRNA regulation [82] and post-translational modifications [83] of proteins have been shown to be important in the context of NAFLD. Despite these limitations, this analysis strongly suggests the role of phenotypic plasticity in the development and progression of NAFLD. It also offers a mechanistic basis for the possible existence of an adipocyte-like phenotype of hepatocytes during NAFLD, where cells may maintain a delicate balance of cellular identity.

## ACKNOWLEDGMENT

MKJ is supported by Ramanujan Fellowship (SB/S2/RJN 049/2018) awarded by the Science and Engineering Research Board (SERB), Department of Science and Technology (DST), Government of India. SS and DS are supported by KVPY fellowship awarded by DST, Government of India. Mr. Atchuta Srinivas Duddu is acknowledged for artwork (Fig 7). The authors would like to thank Mr. Burhanuddin Sabuwala for useful discussions.

## MATERIALS AND METHODS

### a) RAndom CIrcuit PErturbation (RACIPE) analysis

#### 1. RACIPE simulations

RACIPE is a computational method to discern the robust dynamical properties of a particular gene regulatory network topology. It takes in a network topology file as an input and then samples 10000 different sets of parameters. For each parameter set, RACIPE chooses a random set of initial conditions for each node in the network, and solves using Eulers’ method the set of coupled ordinary differential equations (ODEs) that represent the interactions among the different nodes in a network. For each given parameter set and initial conditions, RACIPE reports the steady-state values for each of the nodes in the network.

The parameters are sampled randomly from a specified predefined parameter range for the set of ODEs. We ran RACIPE on our core gene regulatory network shown in Fig 1, using the default parameters given in RACIPE algorithm, but limiting the maximum number of states that are possible to 4. The reason for setting this limit to 4 was that only 7.58 ± 0.08 % of parameter sets enabled > 4 states from the initial analysis using RACIPE.

Note that since we are considering one variable for each node simulated without treating the mRNA and protein levels separately, we name the genes in all caps to represent this dummy variable throughout the study (HNF4A for HNF4α; HNF1A for HNF1α; PPARG for PPARγ and SREBF1 for SREBP-1c). The RACIPE results shown for this network remain largely invariant irrespective of the integration method (Euler vs. Runge-Kutta)/number of initial conditions (n= 1000, 10000, 10000) chosen for every parameter set ***(FIG S6B-D)***.

#### 2. z-score normalizations of the steady state values

RACIPE provides the steady state expression values in log2 scale. We performed z-score normalizations on steady state values that were obtained from the RACIPE simulations so that the distributions/expression values from various nodes could be compared to one another. To compute the z-score transformed expression value for each steady state value (*E*_*i*_) of a given node, we first collated all the untransformed expression values for that node, n. We then computed the mean (*Ē*_*n*_) and the standard deviation (*σ*_*n*_) for this list of untransformed expression values. Then each of the steady state values was transformed by the following formula:

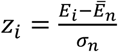

where, *z*_*i*_ is the z-normalized expression value. On plotting the distributions, we found that it is always bimodal and the corresponding z-score value at the central minima of the distribution could segregate the values into two groups which we termed as High (H) and Low (L).

#### 3. Density plots, Bimodality coefficients and clustering analysis

For each node, we plotted the distribution of the z-normalized expression values as a Kernel Density Estimate (KDE) curve. This produces a smoothened curve for the distribution as a probability density function for a finite sample size. We also computed Sarle’s bimodality [84] coefficient for finite samples using the following formula:

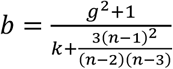

Where n is the number of items in the sample, g is the sample skewness and k is the sample excess kurtosis. As the value of b for the uniform distribution is 5/9, values greater than 5/9 indicate a multimodal distribution (in our case bimodal as is visible from the plots). We performed Unsupervised Hierarchical clustering of the heatmaps which also largely yielded 4 clusters. For clustering analysis for the scatter plots of a pair of genes, we used Hierarchical Agglomerative clustering by fixing the number of clusters that are possible to 4 (H or L for each of the two nodes, hence a maximum of 4 possible clusters).

### b) Relative Stability analysis

To perform a relative stability analysis of a given phase, i.e. coexisting combination of phenotypes such as { HL, LH}, { HL, LH, LL} etc., we collected all the parameters sets that produced that particular phase from the RACIPE analysis. We then explicitly simulated the set of ODEs used in RACIPE in MATLAB for 1000 different initial conditions sampled from a uniform log2 scale. We plotted a Kernel Density Estimate for the distribution of the fraction of the cases for which a particular state was achieved for the given phase.

### c) Dynamic Simulations

#### 1. Bifurcation Diagrams

To plot the bifurcation diagram to show a switch in the levels of HNF4A and PPARG, we used the MATLAB based software MATCONT [85] to simulate the gene regulatory network using a set of Ordinary Differential Equations (ODEs). We used the following set of differential equations – where the rate of change of the expression levels of each node has two terms: Production term (also includes the regulation of that particular node by other nodes) and a Degradation term (assumes first order kinetics). Each interaction (regulation term) in the gene regulatory network between a pair of nodes is represented by a shifted Hills Function (H_S_), where H_S_ for an interaction of B affecting the production of A is defined as [86]:

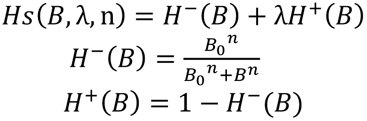

where B_o_ is the threshold value for that interaction, n is the cooperativity for that interaction and λ is the fold change from the basal synthesis rate of A due to B. Therefore, λ > 1 for activators and λ < 1 for inhibitors.

The set of equations that were used to simulate the core gene regulatory network are as follows (green -- maximal production rate; cyan -- regulation term; yellow -- degradation term):

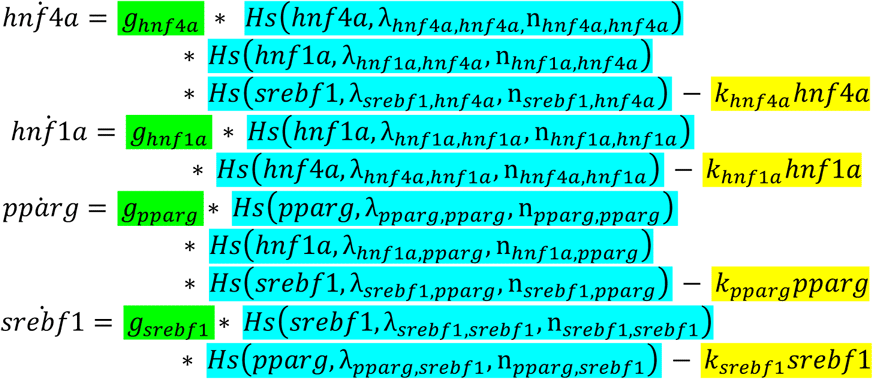

We used the degradation rate of PPARG as the bifurcation parameter. Refer to the **parameter table** for the parameter values used for the simulations.

#### 2. Switching of states

We simulated the set of the above equations explicitly in MATLAB using the ode45 solver and tracked if the levels of PPARG and HNF4A switch over a given time course. We added an additional noise term to each of the equations. The noise term was a random number generated from a Gaussian distribution and multiplied by an amplitude/scaling factor. The amplitude of noise for HNF4A, HNF1A and SREBF1 were kept at 0.1 mimicking the intrinsic noise that is present in the system and 15 for PPARG mimicking the combined effect of intrinsic noise and the extrinsic noise due to a burden of fatty acids to the cells. The following coupled differential equations were used to simulate the switch between the states due to the presence of intrinsic and extrinsic noise. All the parameters, that were used here were the same as those used to create the bifurcation plots. Refer to the table for the parameter values used in these simulations.

The set of equations that were used to simulate the core gene regulatory networks are as follows (green -- maximal production rate; cyan -- regulation term; yellow -- degradation term; red – noise term):

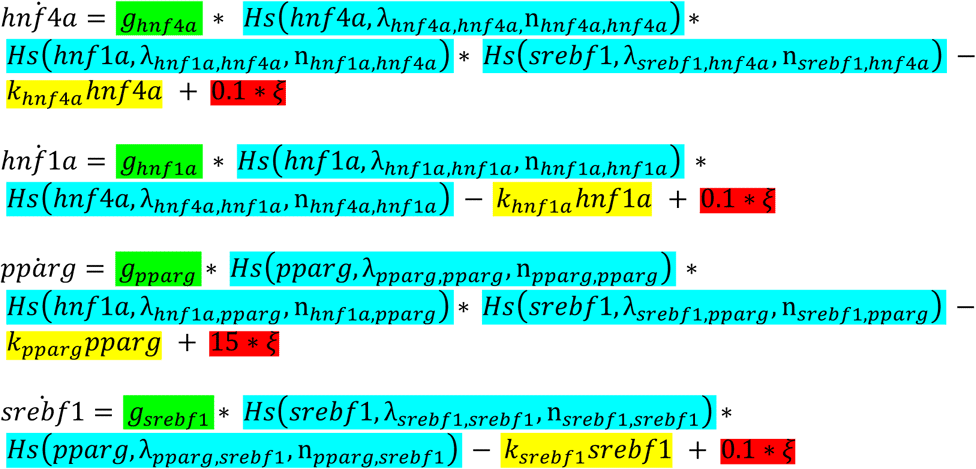

where *ξ* is a random number picked from a normal distribution with a mean of 0 and a standard deviation of 1 and the presence of noise in the system.

### c) Randomization of networks

We created an ensemble of all randomized networks possible using the following rules: for each node, in each instance of randomization of the wild type network ***(FIG 1)***, the indegree and the outdegree of the network was kept fixed. The number of activation edges and the number of inhibitory edges were also kept fixed at 8 and 2 respectively (The same number as that in the wild type network ***(FIG 1)***). Furthermore, the source node and the target node for each of the edges were kept fixed but the identity of the edge in terms of it being an activation or inhibition link was allowed to change. Hence 44 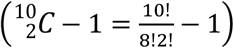 such randomized networks were constructed excluding the wild type case.

### d) Jensen-Shannon Divergence (JSD) and Plasticity scores

For each of the randomized and the wild type network, we calculated the Jensen-Shannon Divergence (JSD) score [57] as follows. We first simulated each of the randomized networks along with the wild type network via RACIPE to obtain the steady state solutions that are possible on a set of 10000 randomly chosen parameter sets. We performed z-score normalizations on the obtained steady state solutions and binarized the expression levels of each of the four genes as High(H) or Low(L) as mentioned in the methods part of RAndom CIrcuit PErturbation (RACIPE) analysis. We then constructed a frequency distribution of all the possible states across mono-stable and multi-stable parameter sets (state frequency distribution) and compared each of the distribution to the reference distribution of the wild-type network to get a corresponding JSD score. All possible 16 (=2^4^) states, emerging from considering the levels of all four nodes in the network, were chosen to calculate the JSD.

In short, for any two discrete frequency distribution P(x) and Q(x),

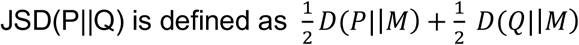

where 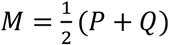

and D stands for the Kullback- Liebler divergence and

is defined as 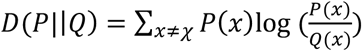

JSD varies from 0 to 1 where 0 corresponds to identical distribution whereas 1 corresponds to no overlap between the distributions ***(FIG S6A)***. This implies that smaller the JSD, the more likely that the state frequency distribution of the randomized network is similar to the wild type network.

The plasticity score (a proxy for the level of plasticity enabled by a gene regulatory network) was defined as:

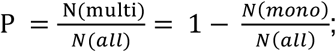

where

N (multi) = Number of parameter sets enabling multi-stable solutions;

N (mono) = Number of parameter sets enabling mono-stable solutions and N (all) = Total number of parameter sets considered

The number of parameters sets that enabled mono-stability or multi-stability were obtain form RACIPE analysis carried on the circuit of interest.

### e) Clinical data analysis

We obtained publicly available transcriptomic data for healthy controls and patients suffering from NAFLD/NASH. We downloaded the preprocessed transcriptomic data from each study and log2 normalized the expression levels if they were not normalized already. The expression values were then used to plot the correlation plots or for comparison of levels of the different genes.

### f) Statistical tests and correlation coefficients

We computed the Spearman correlation coefficients and used the corresponding p-values to gauge the strength of correlation in their expression values between a pair of genes. For statistical comparison values used throughout the study, we used a two-tailed Student’s t-test and computed its significance.

**Table 1:** Production rates and degradation rates of different species:

| Species | Production Rate ($10^6$ molecules $\text{hr}^{-1}$) | Degradation Rate ($\text{hr}^{-1}$) | References |
| --- | --- | --- | --- |
| **HNF4A** | 0.4540 | 0.0491 | [87] |
| **HNF1A** | 0.0570 | 0.0494 | [88] |
| **PPARG** | 0.0628 | 0.08 | [89] |
| **SREBF1** | 1.1058 | 0.1598 | [90] |

The degradation rates were calculated from the reported half-life values of the corresponding proteins assuming first order kinetics i.e. Degradation rate = $\ln 2 / t_{1/2}$ . The production rates were estimated based on typical number of molecules reported in a mammalian cell [86].

**Table 2:**
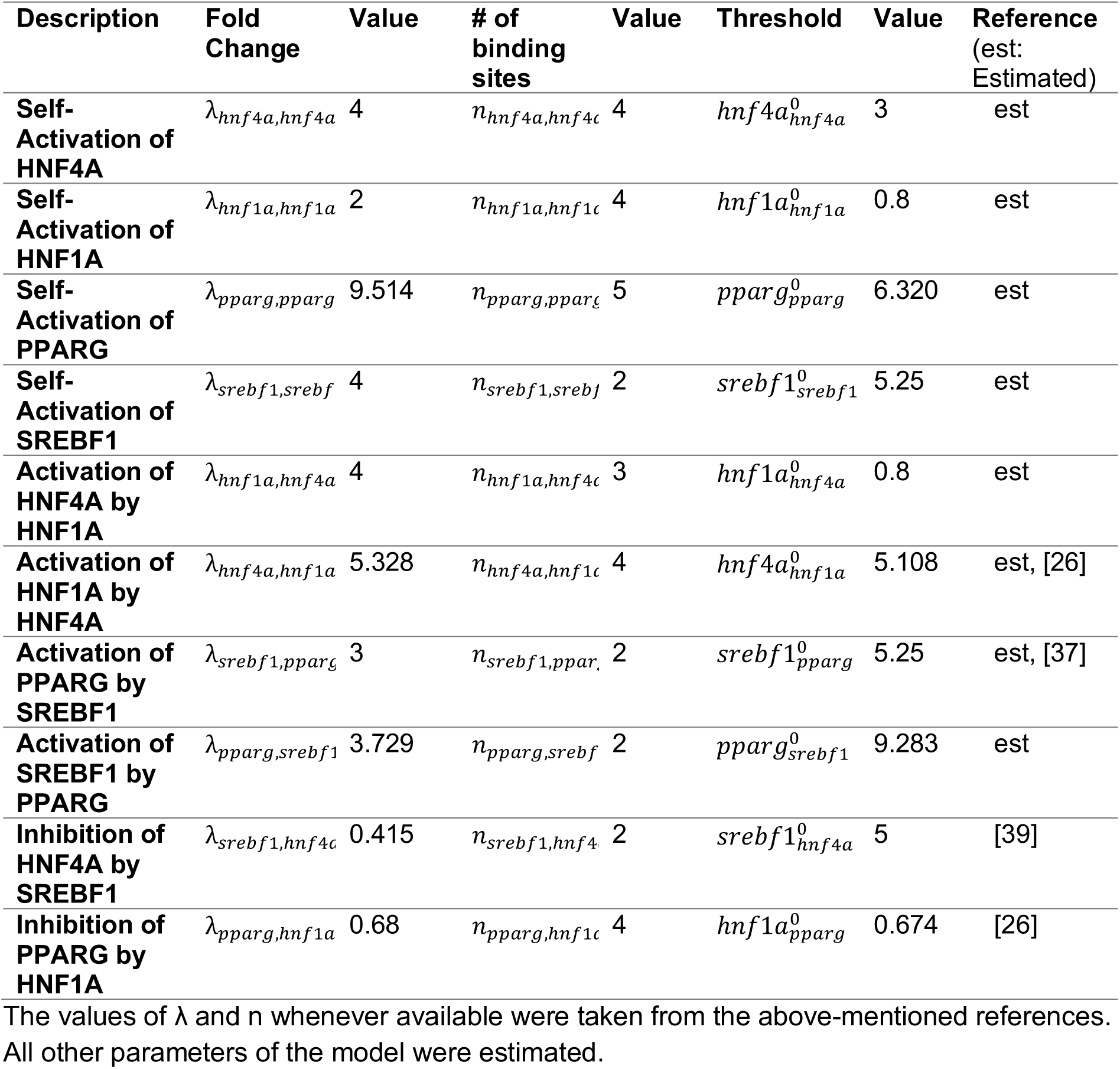
Activation or Inhibition Parameters for the interaction:

## AUTHOR CONTRIBUTIONS

Conceptualization, Mohit Kumar Jolly;

Formal analysis, Sarthak Sahoo and Divyoj Singh; Funding acquisition, Mohit Kumar Jolly;

Methodology, Sarthak Sahoo, Divyoj Singh and Priyanka Chakraborty;

Supervision, Mohit Kumar Jolly;

Writing – original draft, Sarthak Sahoo and Divyoj Singh;

Writing – review & editing, Mohit Kumar Jolly.

## SUPPLEMENTARY FIGURES

**FIGURE S1:**
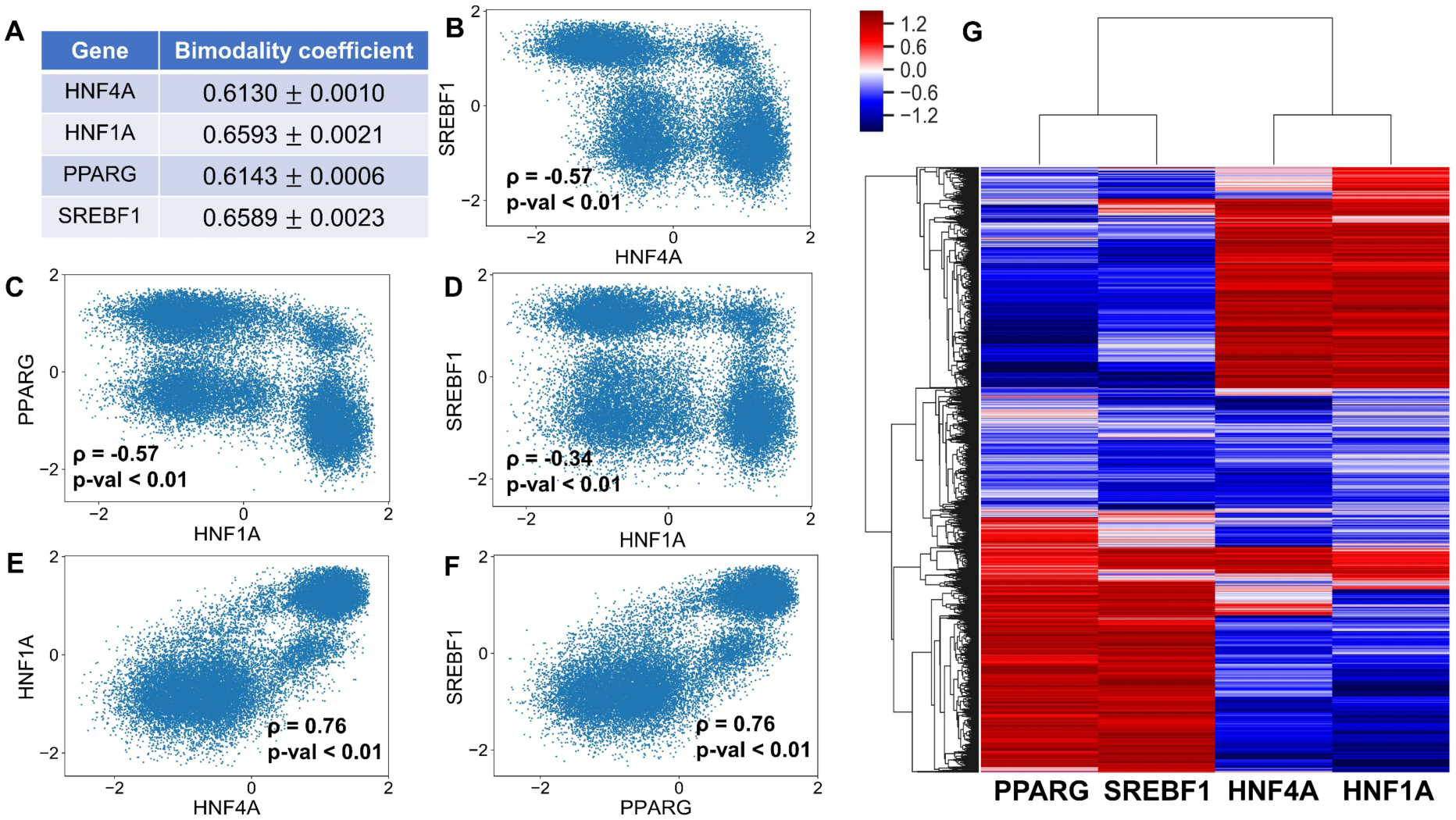
**(A)** Table listing the bimodality coefficients of the four nodes in the core regulatory network over three independent replicates. Mean and standard deviations reflect the statistics over three independent RACIPE replicates. **(B-F)** Scatter plot showing the relationship between HNF4A-SREBF1, HNF1A-PPARG, HNF4A-HNF1A and PPARG-SREBF1 (z-normalized log2 expression values). Spearman’s correlation was performed to obtain the correlation coefficient (ρ) and the significance values (p-val). **(G)** Heatmap showing the relative levels of all 4 nodes of the core gene regulatory network. The color bar represents the relative levels of the individual components (z-normalized log2 expression values).

**FIGURE S2:**
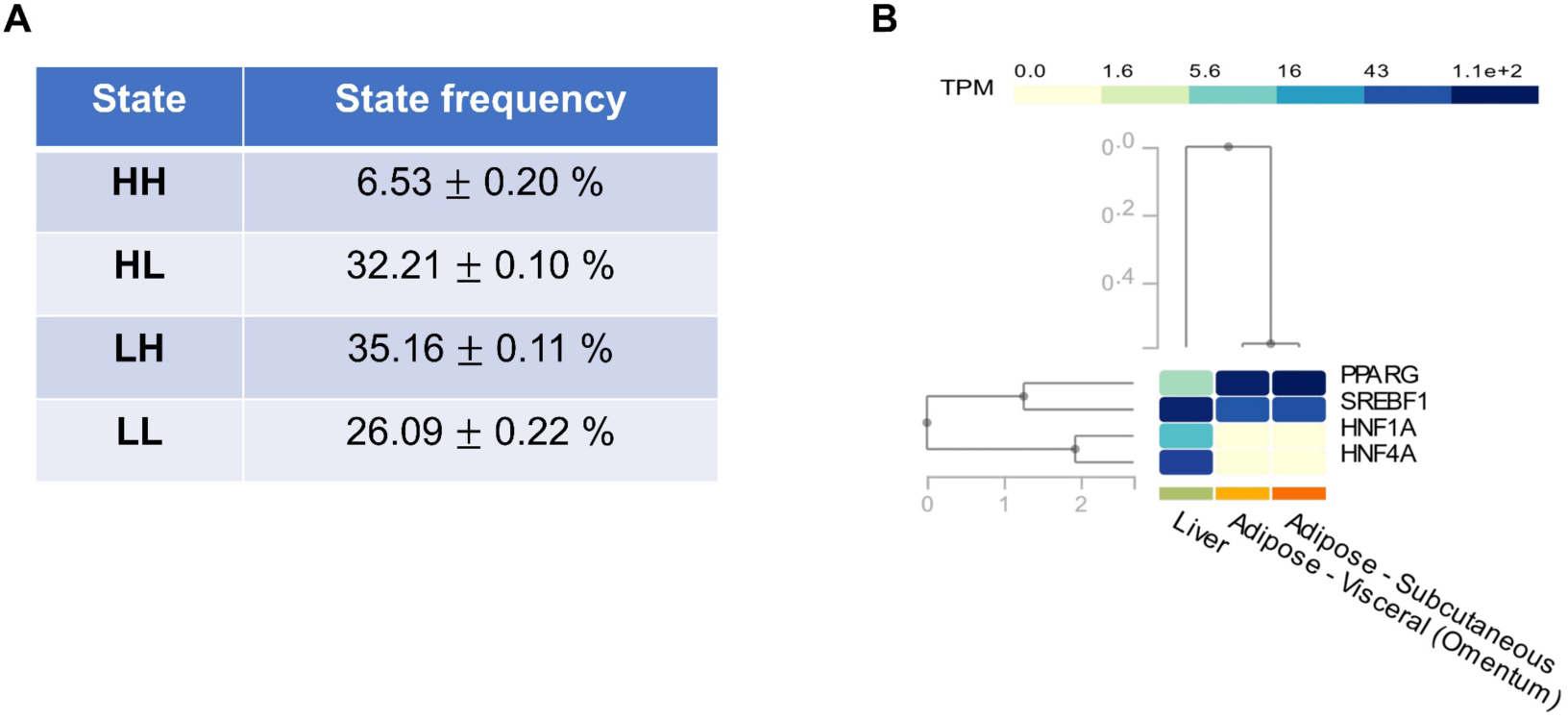
**(A)** Table listing the mean percentage of cases that enable the mentioned state along with their standard deviations (mean ± std. dev) across all the possible phases over three independent RACIPE replicates (n=3; each iteration includes 10000 parameter sets). **(B)** GTEx (Genotype-Tissue Expression) data showing the mRNA expression levels in TPM (Transcripts per million) of the 4 genes of the core regulatory network in Liver and Adipose tissues (Visceral and Subcutaneous) - Data available from GTEx portal on Feb 17, 2020.

**FIGURE S3:**
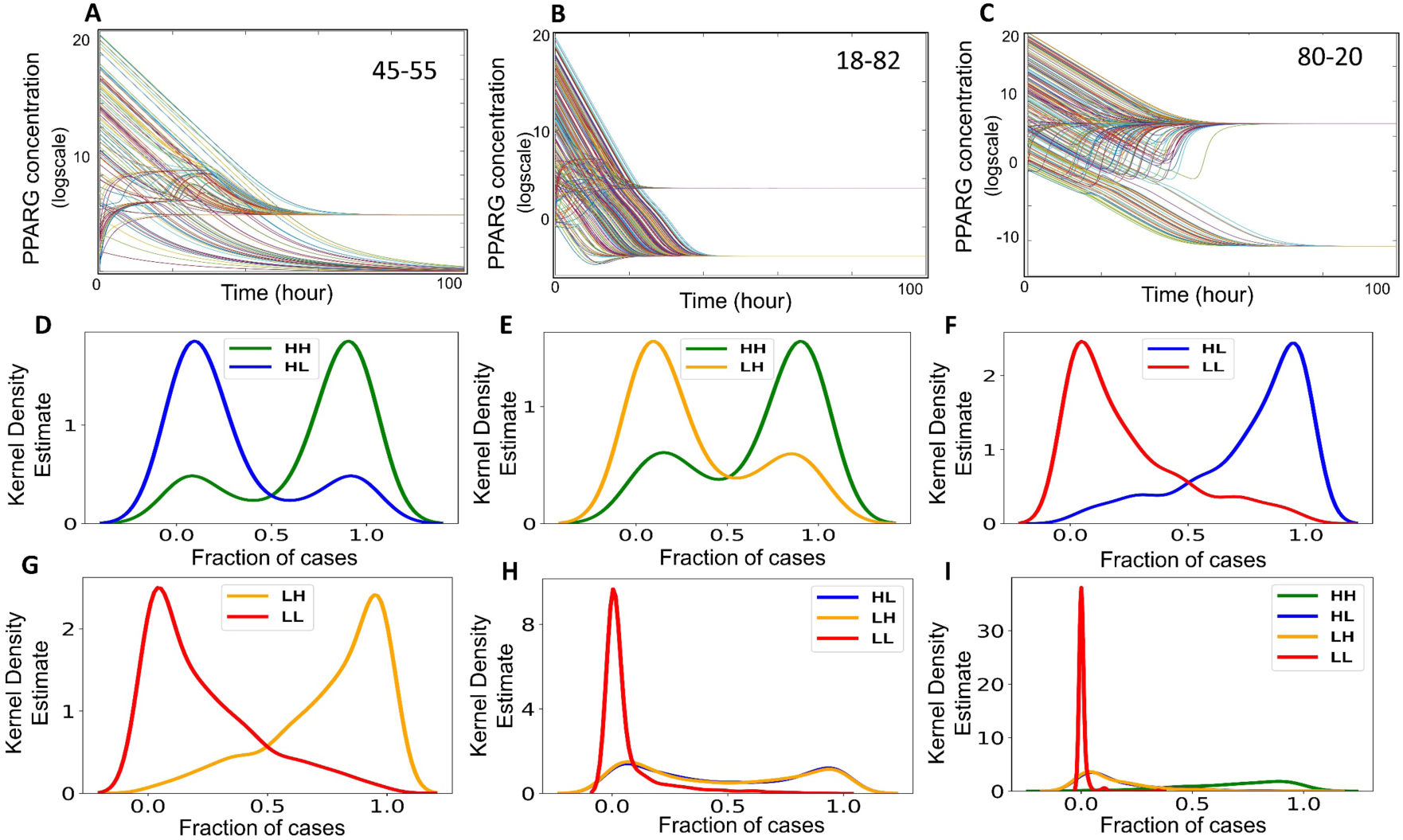
**(A-C)** Schematic representation of relative stability: A system of coupled ordinary differential equations, when simulated for different parameter sets each enabling bistability (two states), can lead to different proportions of initial conditions converging to the two steady states. **(A)** 45 initial conditions lead to state 1, 55 of them lead to state 2. **(B)**18 initial conditions lead to state 1, 82 of them lead to state 2. **(C)** 80 initial conditions lead to state 1, 20 of them lead to state 2. **(D-I)** Kernel density estimates for relative stability of different states in the various multi-stable phases: **(D)** { HH, LH}, **(E)** { HH, HL}, **(F)** { HH, LL}, **(G)** { LH, LL}, **(H)** { HL, LH, LL}, **(I)** { HL, LH, HH, LL}

**FIGURE S4:**
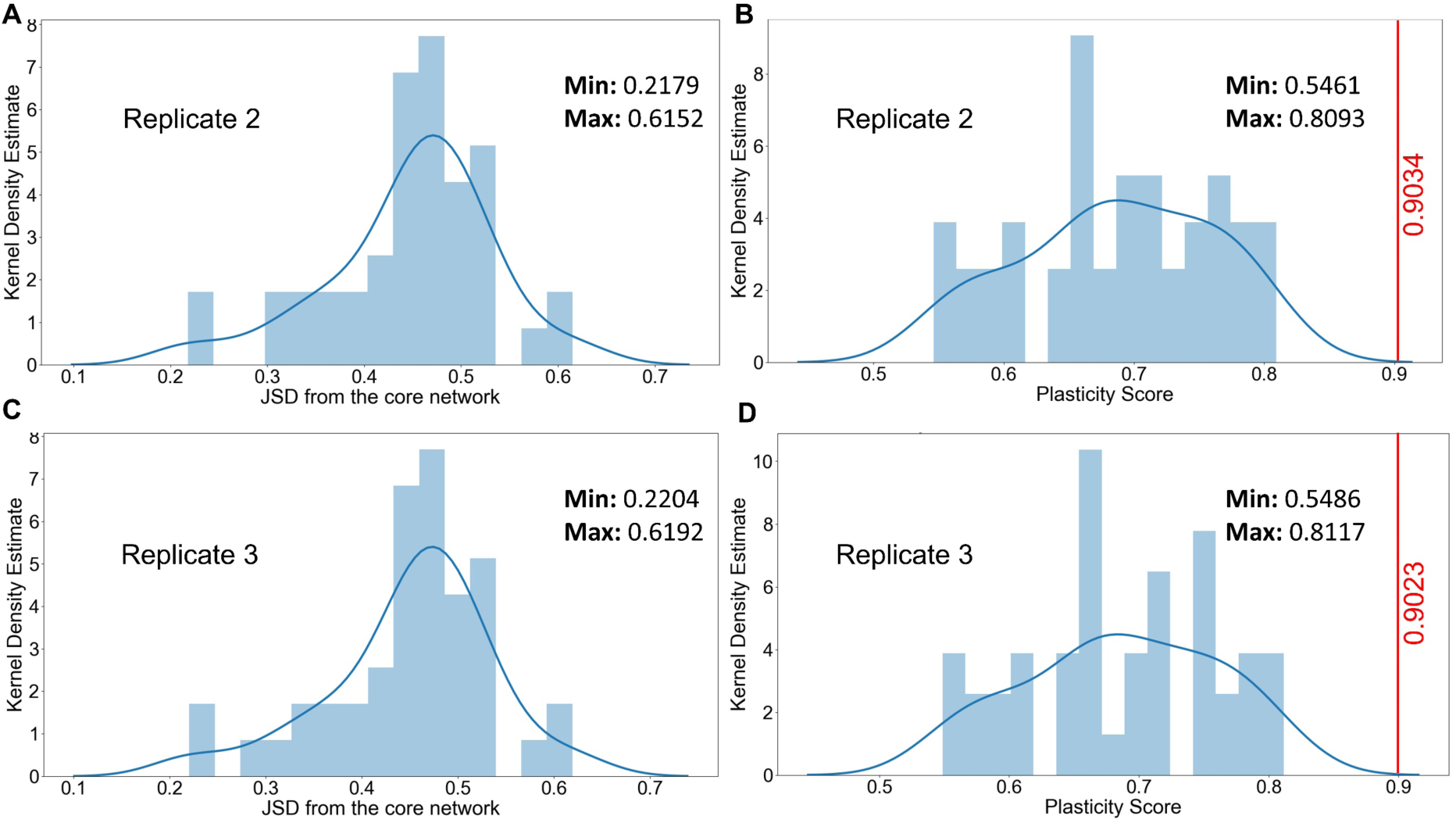
Kernel density estimate plots for Jensen-Shannon Distance (JSD) (A, C) and Plasticity Scores (B, D) for two more RACIPE replicates apart from the one in the main figure 5B-C. In B, D; the red vertical line indicates the plasticity score (=0.90) of the “wild type” core regulatory circuit which is greater than all other “hypothetical” randomized networks.

**FIGURE S5:**
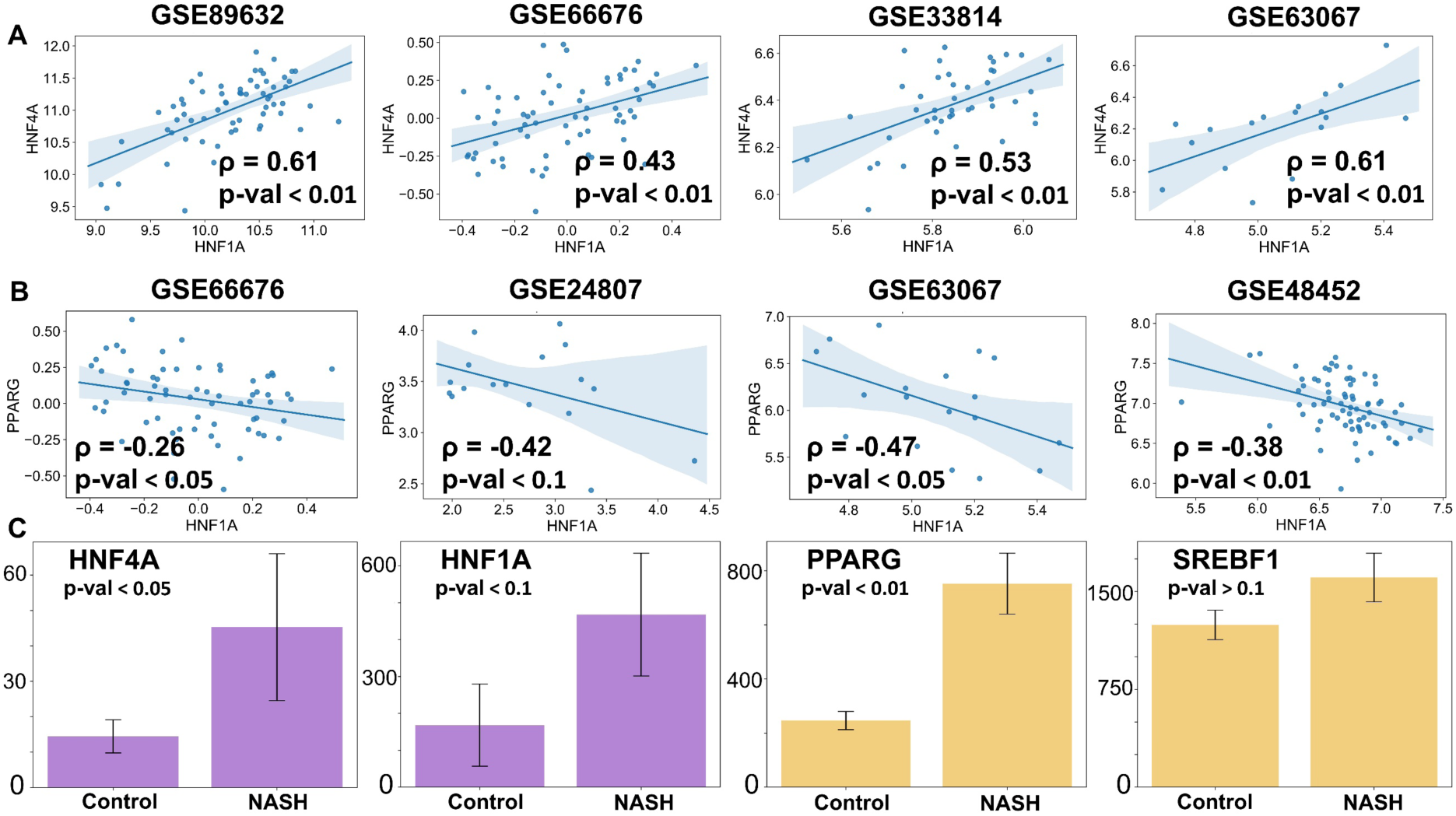
**(A)** Scatter plots between HNF1A and HNF4A in clinical samples. **(B)** Scatter plots between PPARG and SREBF1 in clinical samples. **(C)** Comparison of levels of HNF4A, HNF1A, PPARG and SREBF1 in liver sinusoid endothelial cells for control case vs. mouse model of NASH based on RNA-Seq data. Expression values have been listed as TPM (transcripts per million) for given RNA-seq data. Spearman correlation coefficient is given by ρ, and p-val denotes the corresponding p-value wherever mentioned.

**Figure S6:**
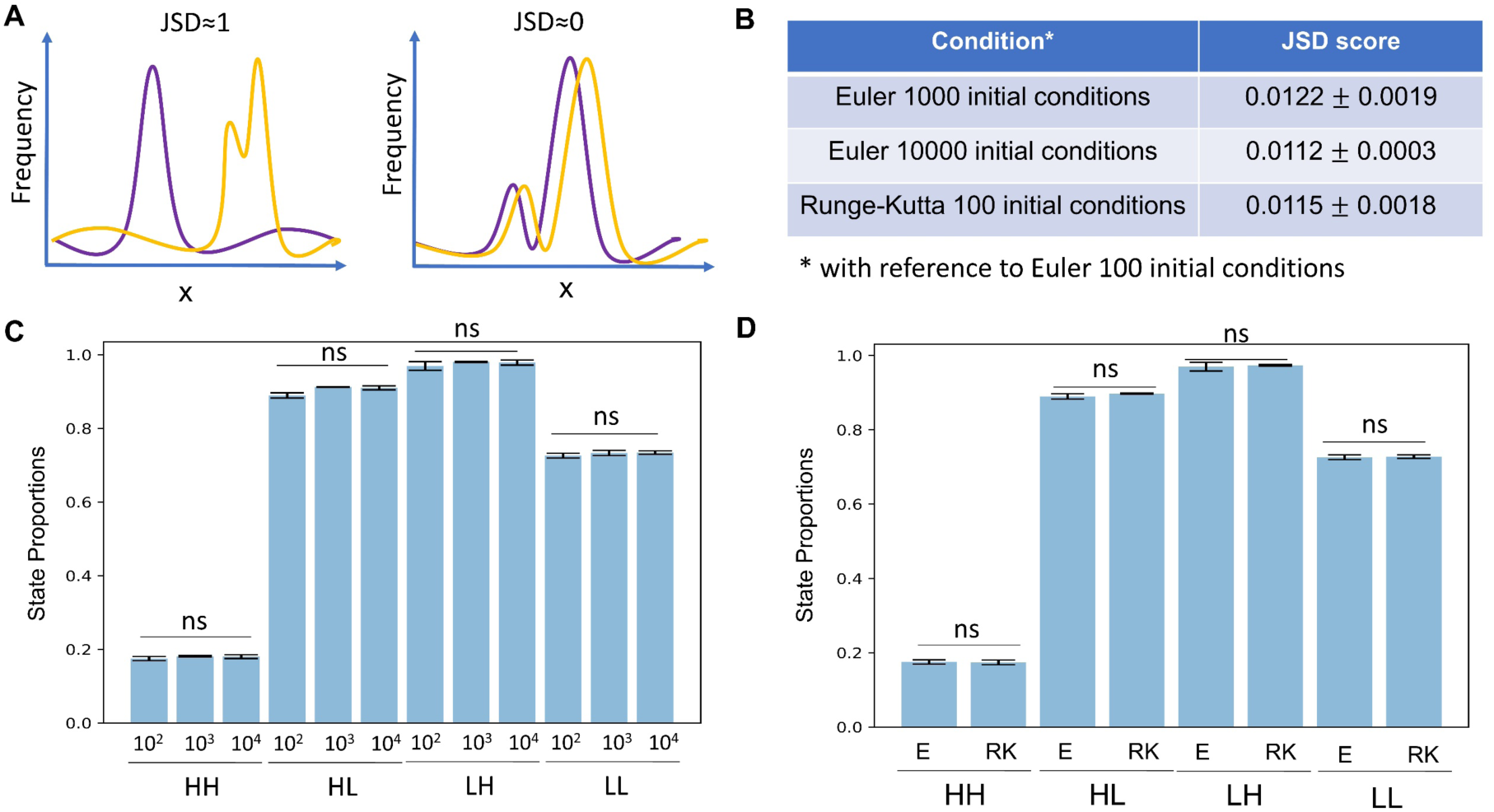
Robustness of methods: **(A)** Schematic describing JSD. **(B)** JSD scores between the frequency of states obtained by different analysis methods. Euler integration method with 100 initial conditions – the default setting for RACIPE – is considered as reference distribution and is used in the rest of this study. **(C)** States proportions of different states do not change significantly if the analysis is done with 100, 1000 or 10000 initial conditions. **(D)** State proportions do not change significantly whether E (Euler integration method) or RK (Runge-Kutta Integration method) is used. ns represents non-significant differences between the different simulation conditions, for students’ t-test done between corresponding 3 replicates.

## REFERENCES

1. Younossi, Z.; Anstee, Q. M.; Marietti, M.; Hardy, T.; Henry, L.; Eslam, M.; George, J.; Bugianesi, E. Global burden of NAFLD and NASH: Trends, predictions, risk factors and prevention. Nat. Rev. Gastroenterol. Hepatol. 2018, 15, 11–20.

2. Abd El-Kader, S. M.; El-Den Ashmawy, E. M. S. Non-alcoholic fatty liver disease: The diagnosis and management. World J. Hepatol. 2015, 7, 846–858.

3. Eslam, M.; Valenti, L.; Romeo, S. Genetics and epigenetics of NAFLD and NASH: Clinical impact. J. Hepatol. 2018, 68, 268–279.

4. Kumar, A.; Shalimar; Walia, G. K.; Gupta, V.; Sachdeva, M. P. Genetics of nonalcoholic fatty liver disease in Asian populations. J. Genet. 2019, 98, 29.

5. Chandrasekharan, K.; Alazawi, W. Genetics of Non-Alcoholic Fatty Liver and Cardiovascular Disease: Implications for Therapy? Front. Pharmacol. 2020, 10, 1413.

6. Eslam, M.; George, J. Genetic contributions to NAFLD: leveraging shared genetics to uncover systems biology. Nat. Rev. Gastroenterol. Hepatol. 2020, 17, 40–52.

7. Arab, J. P.; Arrese, M.; Trauner, M. Recent Insights into the Pathogenesis of Nonalcoholic Fatty Liver Disease. Annu. Rev. Pathol. Mech. Dis. 2018, 13, 321–350.

8. Eslam, M.; Sanyal, A. J.; George, J. MAFLD: A consensus-driven proposed nomenclature for metabolic associated fatty liver disease. Gastroenterology 2020, in press.

9. Sarin, S. K.; Kumar, M.; Eslam, M.; George, J.; Al Mahtab, M.; Akbar, S. M. F.; Jia, J.; Tian, Q.; Aggarwal, R.; Muljono, D. H.; Omata, M.; Ooka, Y.; Han, K. H.; Lee, H. W.; Jafri, W.; Butt, A. S.; Chong, C. H.; Lim, S. G.; Pwu, R. F.; Chen, D. S. Liver diseases in the Asia-Pacific region: a Lancet Gastroenterology & Hepatology Commission. Lancet Gastroenterol. Hepatol. 2020, 5, 167–228.

10. Chen, F.; Esmaili, S.; Rogers, G. B.; Bugianesi, E.; Petta, S.; Marchesini, G.; Bayoumi, A.; Metwally, M.; Azardaryany, M. K.; Coulter, S.; Choo, J. M.; Younes, R.; Rosso, C.; Liddle, C.; Adams, L. A.; Craxì, A.; George, J.; Eslam, M. Lean NAFLD: A Distinct Entity Shaped by Differential Metabolic Adaptation. Hepatology 2019, in press.

11. Duarte, S. M. B.; Stefano, J. T.; Miele, L.; Ponziani, F. R.; Souza-Basqueira, M.; Okada, L. S. R. R.; de Barros Costa, F. G.; Toda, K.; Mazo, D. F. C.; Sabino, E. C.; Carrilho, F. J.; Gasbarrini, A.; Oliveira, C. P. Gut microbiome composition in lean patients with NASH is associated with liver damage independent of caloric intake: A prospective pilot study. Nutr. Metab. Cardiovasc. Dis. 2018, 28, 369–384.

12. Yasutake, K.; Nakamuta, M.; Shima, Y.; Ohyama, A.; Masuda, K.; Haruta, N.; Fujino, T.; Aoyagi, Y.; Fukuizumi, K.; Yoshimoto, T.; Takemoto, R.; Miyahara, T.; Harada, N.; Hayata, F.; Nakashima, M.; Enjoji, M. Nutritional investigation of non-obese patients with non-alcoholic fatty liver disease: The significance of dietary cholesterol. Scand. J. Gastroenterol. 2009, 44, 471–477.

13. Zeng, L.; Tang, W. J.; Yin, J. J.; Zhou, B. J. Signal transductions and nonalcoholic fatty liver: a mini-review. Int J Clin Exp Med 2014, 7, 1624–1631.

14. Wafer, R.; Tandon, P.; Minchin, J. E. N. The role of peroxisome proliferator-activated receptor gamma (PPARG) in adipogenesis: Applying knowledge from the fish aquaculture industry to biomedical research. Front. Endocrinol. (Lausanne). 2017, 8, 102.

15. Lee, Y. K.; Park, J. E.; Lee, M.; Hardwick, J. P. Hepatic lipid homeostasis by peroxisome proliferator-activated receptor gamma 2. Liver Res. 2018, 2, 209–215.

16. Lake, A. D.; Chaput, A. L.; Novak, P.; Cherrington, N. J.; Smith, C. L. Transcription factor binding site enrichment analysis predicts drivers of altered gene expression in nonalcoholic steatohepatitis. Biochem. Pharmacol. 2016, 122, 62–71.

17. Baciu, C.; Pasini, E.; Angeli, M.; Schwenger, K.; Afrin, J.; Humar, A.; Fischer, S.; Patel, K.; Allard, J.; Bhat, M. Systematic integrative analysis of gene expression identifies HNF4A as the central gene in pathogenesis of non-alcoholic steatohepatitis. PLoS One 2017, 12, e0189223.

18. Lau, H. H.; Ng, N. H. J.; Loo, L. S. W.; Jasmen, J. B.; Teo, A. K. K. The molecular functions of hepatocyte nuclear factors – In and beyond the liver. J. Hepatol. 2018, 68, 1033–1048.

19. Bonzo, J. A.; Ferry, C. H.; Matsubara, T.; Kim, J. H.; Gonzalez, F. J. Suppression of hepatocyte proliferation by hepatocyte nuclear factor 4α in adult mice. J. Biol. Chem. 2012, 287, 7345–56.

20. Huck, I.; Gunewardena, S.; Espanol-Suner, R.; Willenbring, H.; Apte, U. Hepatocyte Nuclear Factor 4 Alpha Activation Is Essential for Termination of Liver Regeneration in Mice. Hepatology 2019, 70, 666–681.

21. Ni, Q.; Ding, K.; Wang, K. Q.; He, J.; Yin, C.; Shi, J.; Zhang, X.; Xie, W. F.; Shi, Y. Q. Deletion of HNF1α in hepatocytes results in fatty liver-related hepatocellular carcinoma in mice. FEBS Lett. 2017, 591, 1947–1957.

22. Odom, D. T.; Dowell, R. D.; Jacobsen, E. S.; Nekludova, L.; Rolfe, P. A.; Danford, T. W.; Gifford, D. K.; Fraenkel, E.; Bell, G. I.; Young, R. A. Core transcriptional regulatory circuitry in human hepatocytes. Mol. Syst. Biol. 2006, 2, 2006.0017.

23. Li, J.; Ning, G.; Duncan, S. A. Mammalian hepatocyte differentiation requires the transcription factor HNF-4α. Genes Dev. 2000.

24. Hayhurst, G. P.; Lee, Y.-H.; Lambert, G.; Ward, J. M.; Gonzalez, F. J. Hepatocyte Nuclear Factor 4 (Nuclear Receptor 2A1) Is Essential for Maintenance of Hepatic Gene Expression and Lipid Homeostasis. Mol. Cell. Biol. 2001.

25. Martinez-Jimenez, C. P.; Kyrmizi, I.; Cardot, P.; Gonzalez, F. J.; Talianidis, I. Hepatocyte Nuclear Factor 4 Coordinates a Transcription Factor Network Regulating Hepatic Fatty Acid Metabolism. Mol. Cell. Biol. 2010, 30, 565–577.

26. Patitucci, C.; Couchy, G.; Bagattin, A.; Cañeque, T.; De Reyniès, A.; Scoazec, J. Y.; Rodriguez, R.; Pontoglio, M.; Zucman-Rossi, J.; Pende, M.; Panasyuk, G. Hepatocyte nuclear factor 1α suppresses steatosisassociated liver cancer by inhibiting PPARγ transcription. J. Clin. Invest. 2017, 127, 1873–1888.

27. Tontonoz, P.; Hu, E.; Spiegelman, B. M. Stimulation of adipogenesis in fibroblasts by PPARγ2, a lipid-activated transcription factor. Cell 1994.

28. Softic, S.; Cohen, D. E.; Kahn, C. R. Role of Dietary Fructose and Hepatic De Novo Lipogenesis in Fatty Liver Disease. Dig. Dis. Sci. 2016, 61, 1282–93.

29. Lambert, J. E.; Ramos-Roman, M. A.; Browning, J. D.; Parks, E. J. Increased de novo lipogenesis is a distinct characteristic of individuals with nonalcoholic fatty liver disease. Gastroenterology 2014, 146, 726–35.

30. Kim, J. B.; Spiegelman, B. M. ADD1/SREBP1 promotes adipocyte differentiation and gene expression linked to fatty acid metabolism. Genes Dev. 1996, 10, 1096–107.

31. Pettinelli, P.; Videla, L. A. Up-regulation of PPAR-γ mRNA expression in the liver of obese patients: An additional reinforcing lipogenic mechanism to SREBP-1c induction. J. Clin. Endocrinol. Metab. 2011, 96, 1424–30.

32. Kim, D. H.; Kim, J.; Kwon, J. S.; Sandhu, J.; Tontonoz, P.; Lee, S. K.; Lee, S.; Lee, J. W. Critical Roles of the Histone Methyltransferase MLL4/KMT2D in Murine Hepatic Steatosis Directed by ABL1 and PPARγ2. Cell Rep. 2016, 17, 1671–1682.

33. Bahrami-Nejad, Z.; Zhao, M.; Tholen, S.; Hunerdosse, D.; Tkach, K.; van Schie, S.; Chung, M.; Teruel, M. A Transcriptional Circuit Filters Oscillating Circadian Hormonal Inputs to Regulate Fat Cell Differentiation. Cell Metab. 2018, 27, 854–868.

34. Shao, W.; Espenshade, P. J. Expanding roles for SREBP in metabolism. Cell Metab. 2012, 16, 414–419.

35. Zhang, L.; Li, C.; Wang, F.; Zhou, S.; Shangguan, M.; Xue, L.; Zhang, B.; Ding, F.; Hui, D.; Liang, A.; He, D. Treatment with PPAR α agonist clofibrate inhibits the transcription and activation of srebps and reduces triglyceride and cholesterol levels in liver of broiler chickens. PPAR Res. 2015, 2015, 347245.

36. Fajas, L.; Schoonjans, K.; Gelman, L.; Kim, J. B.; Najib, J.; Martin, G.; Fruchart, J.-C.; Briggs, M.; Spiegelman, B. M.; Auwerx, J. Regulation of Peroxisome Proliferator-Activated Receptor γ Expression by Adipocyte Differentiation and Determination Factor 1/Sterol Regulatory Element Binding Protein 1: Implications for Adipocyte Differentiation and Metabolism. Mol. Cell. Biol. 1999, 19, 5495–5503.

37. Kim, J. B.; Wright, H. M.; Wright, M.; Spiegelman, B. M. ADD1/SREBP1 activates PPARγ through the production of endogenous ligand. Proc Natl Acad Sci U S A 1998, 95, 4333–4337.

38. Fang, L.; Zhang, M.; Li, Y.; Liu, Y.; Cui, Q.; Wang, N. PPARgene: A Database of Experimentally Verified and Computationally Predicted PPAR Target Genes. PPAR Res. 2016, 2016, 6042162.

39. Xie, X.; Liao, H.; Dang, H.; Pang, W.; Guan, Y.; Wang, X.; Shyy, J. Y. J.; Zhu, Y.; Sladek, F. M. Down-regulation of Hepatic HNF4α Gene Expression during Hyperinsulinemia via SREBPs. Mol. Endocrinol. 2009, 23, 434–43.

40. Yamamoto, T.; Shimano, H.; Nakagawa, Y.; Ide, T.; Yahagi, N.; Matsuzaka, T.; Nakakuki, M.; Takahashi, A.; Suzuki, H.; Sone, H.; Toyoshima, H.; Sato, R.; Yamada, N. SREBP-1 Interacts with Hepatocyte Nuclear Factor-4α and Interferes with PGC-1 Recruitment to Suppress Hepatic Gluconeogenic Genes. J. Biol. Chem. 2004, 279, 12027–12035.

41. Zhou, J. X.; Huang, S. Understanding gene circuits at cell-fate branch points for rational cell reprogramming. Trends Genet. 2011, 27, 55–62.

42. Jia, D.; Jolly, M. K.; Tripathi, S. C.; Hollander, P. Den; Huang, B.; Lu, M.; Celiktas, M.; Ramirez-Pena, E.; Ben-Jacob, E.; Onuchic, J. N.; Hanash, S. M.; Mani, S. A.; Levine, H. Distinguishing Mechanisms Underlying EMT Tristability. Cancer Converg. 2017, 1, 2.

43. Huang, B.; Lu, M.; Jia, D.; Ben-Jacob, E.; Levine, H.; Onuchic, J. N. Interrogating the topological robustness of gene regulatory circuits by randomization. PLoS Comput. Biol. 2017, 13, e1005456.

44. Waddington, C. H. The strategy of the genes. A discussion of some aspects of theoretical biology. With an appendix by H. Kacser. Strateg. genes. A Discuss. some Asp. Theor. Biol. With an Append. by H. Kacser. 1957.

45. Khurana, S.; Jaiswal, A. K.; Mukhopadhyay, A. Hepatocyte nuclear factor-4α induces transdifferentiation of hematopoietic cells into hepatocytes. J. Biol. Chem. 2010, 285, 4725–4731.

46. Mooney, S. M.; Jolly, M. K.; Levine, H.; Kulkarni, P. Phenotypic plasticity in prostate cancer: role of intrinsically disordered proteins. Asian J. Androl. 2016, 18, 704–10.

47. Huang, S. Hybrid T-Helper Cells: Stabilizing the Moderate Center in a Polarized System. PLoS Biol. 2013, 11, e1001632.

48. Elowitz, M. B.; Levine, A. J.; Siggia, E. D.; Swain, P. S. Stochastic gene expression in a single cell. Science 2002, 297, 1183–6.

49. Tripathi, S.; Levine, H.; Jolly, M. K. A Mechanism for Epithelial-Mesenchymal Heterogeneity in a Population of Cancer Cells. bioRxiv 2019, 592691.

50. Jolly, M. K.; Tripathi, S. C.; Somarelli, J. A.; Hanash, S. M.; Levine, H. Epithelial-mesenchymal plasticity: How have quantitative mathematical models helped improve our understanding ? Mol. Oncol. 2017, 11, 739–754.

51. Huang, B.; Jolly, M. K.; Lu, M.; Tsarfaty, I.; Ben-Jacob, E.; Onuchic, J. N. Modeling the Transitions between Collective and Solitary Migration Phenotypes in Cancer Metastasis. Sci. Rep. 2015, 5.

52. Wheeler, M. C.; Gekakis, N. Hsp90 modulates PPARγ activity in a mouse model of nonalcoholic fatty liver disease. J. Lipid Res. 2014, 55, 1702–1710.

53. Lakhani, H. V.; Sharma, D.; Dodrill, M. W.; Nawab, A.; Sharma, N.; Cottrill, C. L.; Shapiro, J. I.; Sodhi, K. Phenotypic alteration of hepatocytes in non-alcoholic fatty liver disease. Int. J. Med. Sci. 2018, 15, 1591–1599.

54. Carpino, G.; Renzi, A.; Onori, P.; Gaudio, E. Role of hepatic progenitor cells in nonalcoholic fatty liver disease development: Cellular cross-talks and molecular networks. Int. J. Mol. Sci. 2013, 14, 20112–20130.

55. Balázsi, G.; van Oudenaarden, A.; Collins, J. J.; van Oudenaarden, A.; Collins, J. J. Cellular decision making and biological noise: from microbes to mammals. Cell 2011, 144, 910–925.

56. Choudhary, N. S.; Saraf, N.; Saigal, S.; Gautam, D.; Lipi, L.; Rastogi, A.; Goja, S.; Menon, P. B.; Bhangui, P.; Ramchandra, S. K.; Soin, A. S. Rapid Reversal of Liver Steatosis With Life Style Modification in Highly Motivated Liver Donors. J. Clin. Exp. Hepatol. 2015, 5, 123–126.

57. Lin, J. Divergence Measures Based on the Shannon Entropy. IEEE Trans. Inf. Theory 1991, 37.

58. Hari, K.; Sabuwala, B.; Subramani, B. V.; Porta, C. La; Zapperi, S.; Font-Clos, F.; Jolly, M. K. Identifying inhibitors of epithelial-mesenchymal plasticity using a network topology based approach. bioRxiv 2019, 854307.

59. Benedict, M.; Zhang, X. Non-alcoholic fatty liver disease: An expanded review. World J. Hepatol. 2017, 9, 715–732.

60. Del Campo, J. A.; Gallego-Durán, R.; Gallego, P.; Grande, L. Genetic and epigenetic regulation in nonalcoholic fatty liver disease (NAFLD). Int. J. Mol. Sci. 2018, 19.

61. Ma, M.; Duan, R.; Zhong, H.; Liang, T.; Guo, L. The crosstalk between fat homeostasis and liver regional immunity in NAFLD. J. Immunol. Res. 2019, 2019.

62. Blencowe; Karunanayake; Wier; Hsu; Yang Network Modeling Approaches and Applications to Unravelling Non-Alcoholic Fatty Liver Disease. Genes (Basel). 2019, 10, 966.

63. Shubham, K.; Vinay, L.; Vinod, P. K. Systems-level organization of non-alcoholic fatty liver disease progression network. Mol. Biosyst. 2017, 13, 1898–1911.

64. Ashworth, W. B.; Davies, N. A.; Bogle, I. D. L. A Computational Model of Hepatic Energy Metabolism: Understanding Zonated Damage and Steatosis in NAFLD. PLOS Comput. Biol. 2016, 12, e1005105.

65. Maldonado, E. M.; Fisher, C. P.; Mazzatti, D. J.; Barber, A. L.; Tindall, M. J.; Plant, N. J.; Kierzek, A. M.; Moore, J. B. Multi-scale, whole-system models of liver metabolic adaptation to fat and sugar in non-alcoholic fatty liver disease. npj Syst. Biol. Appl. 2018, 4, 1–10.

66. Pirola, C. J.; Sookoian, S. Tackling the complexity of nonalcoholic steatohepatitis treatment: challenges and opportunities based on systems biology and machine learning approaches. HepatoBiliary Surg. Nutr. 2018, 7, 495–498.

67. Eslam, M.; George, J. Genetic Insights for Drug Development in NAFLD. Trends Pharmacol. Sci. 2019, 40, 506–516.

68. Pirola, C. J.; Sookoian, S. Multiomics biomarkers for the prediction of nonalcoholic fatty liver disease severity. World J. Gastroenterol. 2018, 24, 1601–1615.

69. Guantes, R.; Poyatos, J. F. Multistable decision switches for flexible control of epigenetic differentiation. PLoS Comput. Biol. 2008, 4, e1000235.

70. Ozbudak, E. M.; Thattai, M.; Lim, H. N.; Shraiman, B. I.; van Oudenaarden, A. Multistability in the lactose utilization network of Escherichia coli. Nature 2004, 427, 737–40.

71. Jolly, M. K.; Celia-Terrassa, T. Dynamics of Phenotypic Heterogeneity Associated with EMT and Stemness during Cancer Progression. J Clin Med 2019, 8, 1542.

72. Parafati, M.; Kirby, R. J.; Khorasanizadeh, S.; Rastinejad, F.; Malany, S. A nonalcoholic fatty liver disease model in human induced pluripotent stem cell-derived hepatocytes, created by endoplasmic reticulum stress-induced steatosis. DMM Dis. Model. Mech. 2018, 11.

73. Tripathi, S.; Chakraborty, P.; Levine, H.; Jolly, M. K. A mechanism for epithelial-mesenchymal heterogeneity in a population of cancer cells. PLoS Comput Biol 2020, 16, e1007619.

74. Jia, W.; Deshmukh, A.; Mani, S. A.; Jolly, M. K.; Levine, H. A possible role for epigenetic feedback regulation in the dynamics of the Epithelial-Mesenchymal Transition (EMT). Phys. Biol. 2019, 16, 066004.

75. Farrell, G. C.; van Rooyen, D.; Gan, L.; Chitturi, S. NASH is an inflammatory disorder: Pathogenic, prognostic and therapeutic implications. Gut Liver 2012, 6, 149–171.

76. Ganz, M.; Szabo, G. Immune and inflammatory pathways in NASH. Hepatol. Int. 2013, 7, S771–S781.

77. Chattopadhyay, T.; Maniyadath, B.; Bagul, H. P.; Chakraborty, A.; Shukla, N.; Budnar, S.; Rajendran, A.; Shukla, A.; Kamat, S. S.; Kolthur-Seetharam, U. Spatiotemporal gating of SIRT1 functions by O-GlcNAcylation is essential for liver metabolic switching and prevents hyperglycemia. Proc Natl Acad Sci U S A 2020, in press.

78. Horton, J. D.; Bashmakov, Y.; Shimomura, I.; Shimano, H. Regulation of sterol regulatory element binding proteins in livers of fasted and refed mice. Proc. Natl. Acad. Sci. U. S. A. 1998, 95, 5987–5992.

79. Billon, N.; Dani, C. Developmental Origins of the Adipocyte Lineage: New Insights from Genetics and Genomics Studies. Stem Cell Rev. Reports 2012, 8, 55–66.

80. Zaret, K. S. Hepatocyte differentiation: From the endoderm and beyond. Curr. Opin. Genet. Dev. 2001, 11, 568–574.

81. Li, H.; Zhu, L.; Chen, H.; Li, T.; Han, Q.; Wang, S.; Yao, X.; Feng, H.; Fan, L.; Gao, S.; Boyd, R.; Cao, X.; Zhu, P.; Li, J.; Keating, A.; Su, X.; Zhao, R. C. Generation of Functional Hepatocytes from Human Adipose-Derived MYC ^+^ KLF4 ^+^ GMNN ^+^ Stem Cells Analyzed by Single-Cell RNA-Seq Profiling. Stem Cells Transl. Med. 2018, 7, 792–805.

82. Dongiovanni, P.; Meroni, M.; Longo, M.; Fargion, S.; Fracanzani, A. L. MiRNA signature in NAFLD: A turning point for a non-invasive diagnosis. Int. J. Mol. Sci. 2018, 19.

83. Cheng, Y.; Hou, T.; Ping, J.; Chen, G.; Chen, J. Quantitative succinylome analysis in the liver of non-alcoholic fatty liver disease rat model. Proteome Sci. 2016, 14, 3.

84. Ellison, A. M. Effect of Seed Dimorphism on the Density-Dependent Dynamics of Experimental Populations of Atriplex triangularis (Chenopodiaceae). Am. J. Bot. 1987, 74, 1280.

85. Dhooge, A.; Govaerts, W.; Kuznetsov, Y. A.; Meijer, H. G. E.; Sautois, B. New features of the software MatCont for bifurcation analysis of dynamical systems. Math. Comput. Model. Dyn. Syst. 2008, 14, 147–175.

86. Lu, M.; Jolly, M. K.; Levine, H.; Onuchic, J. N.; Ben-Jacob, E. MicroRNA-based regulation of epithelial-hybrid-mesenchymal fate determination. Proc. Natl. Acad. Sci. U. S. A. 2013, 110, 18174–9.

87. Mathieson, T.; Franken, H.; Kosinski, J.; Kurzawa, N.; Zinn, N.; Sweetman, G.; Poeckel, D.; Ratnu, V. S.; Schramm, M.; Becher, I.; Steidel, M.; Noh, K. M.; Bergamini, G.; Beck, M.; Bantscheff, M.; Savitski, M. M. Systematic analysis of protein turnover in primary cells. Nat. Commun. 2018, 9, 1–10.

88. Dong, B.; Li, H.; Singh, A. B.; Cao, A.; Liu, J. Inhibition of PCSK9 transcription by Berberine involves down-regulation of hepatic HNF1α protein expression through the ubiquitin-proteasome degradation pathway. J. Biol. Chem. 2015, 290, 4047–4058.

89. Waite, K. J.; Floyd, Z. E.; Arbour-Reily, P.; Stephens, J. M. Interferon-γ-induced Regulation of Peroxisome Proliferator-activated Receptor γ and STATs in Adipocytes. J. Biol. Chem. 2001, 276, 7062–7068.

90. Hirano, Y.; Yoshida, M.; Shimizu, M.; Sato, R. Direct Demonstration of Rapid Degradation of Nuclear Sterol Regulatory Element-binding Proteins by the Ubiquitin-Proteasome Pathway. J. Biol. Chem. 2001, 276, 36431–36437.

